# Sensory context gates oxytocin effects on auditory processing in macaques

**DOI:** 10.64898/2026.11.10.765620

**Authors:** M. Froesel, H. Wang, W. Vanduffel, Q. Zhu

## Abstract

Oxytocin shapes social perception, yet its neural effects vary substantially across experiments and clinical trials yield inconsistent results. A key unresolved question is whether oxytocin modulates early sensory processing in primates as a function of sensory context. Using high-resolution pharmacological fMRI in two awake rhesus macaques, we compared systemic oxytocin with saline during unimodal auditory and audiovisual tasks. During unimodal auditory stimulation, oxytocin attenuated sound-evoked responses throughout auditory core, belt and associative regions for both vocal and non-vocal sounds, while distributed sound-category information remained decodable. In contrast, when the same vocalizations were presented within an audiovisual task, this attenuation disappeared and responses increased in a restricted set of auditory core and belt regions. Oxytocin also reorganized resting-state coupling among auditory, visual and prefrontal systems and changed how evoked responses covaried across runs. Together, these findings suggest that oxytocin modulates sensory response magnitude and inter-regional coordination in a context-dependent manner rather than uniformly enhancing social signals.

## Introduction

Over the past two decades, oxytocin (OT) has become one of the most intensively studied neuromodulators of social behavior. In humans, intranasal administration has been reported to increase trust (Kosfeld et al., 2005), gaze to the eye region and memory for face identity, and to alter the neural responses within social-processing circuits (Barton et al., 2025; Baumgartner et al., 2008; Guastella et al., 2008; Savaskan et al., 2008). These findings have contributed to the social-salience hypothesis of OT, which proposes that OT increases the salience of socially relevant cues. Yet, its effects are strongly shaped by contextual factors such as individual state and task demands (Bartz et al., 2011; Shamay-Tsoory & Abu-Akel, 2016, Barton et al., 2025). Complementing this account, anxiolytic models emphasize OT’s ability to dampen threat-related and defensive responses, including amygdala responses to fearful stimuli (Kirsch et al., 2005). These two views are not mutually exclusive: oxytocin may enhance, attenuate, or reorganize neural responses depending on the sensory context and the behavioral relevance of incoming information. Yet, neither framework directly addresses whether OT acts at the level of sensory processing itself, and if it does, what determines the direction of this modulation.

This question is particularly pressing for the auditory system. Conspecific vocalizations are central to primate social communication (Cheney & Seyfarth, 2018), yet how oxytocin shapes auditory processing remains poorly understood. In humans, intranasal OT modulates responses to voices and enhance responses in voice-sensitive superior temporal cortex during identity recognition (Borowiak & von Kriegstein, 2020). Evidence from rodents, however, suggests that oxytocin acts at early sensory stages. OT contributes to rodent social recognition through olfactory mechanisms, modulates the excitation–inhibition balance of auditory cortex so that pup calls become salient (Marlin et al., 2015; Ross & Young, 2009), and suppresses spontaneous activity across auditory cortex, medial prefrontal cortex and amygdala while strengthening bottom-up auditory–prefrontal coupling (Jung et al., 2026). Whether a primate brain expresses comparable modulation is an open question, because receptor architecture differs markedly across species. Mice express dense oxytocin receptors in auditory cortex, prefrontal cortex and amygdala (Mitre et al., 2016). Human transcriptomic data suggest relatively enriched oxytocin receptor (OXTR) expression in subcortical and olfactory regions (Quintana et al., 2019) and rhesus macaques show binding largely confined to subcortical structures with little neocortical expression (Freeman et al., 2014). Macaques therefore provide a valuable test of whether sensory and network-level modulation described in rodents can be expressed in a primate brain with a different receptor architecture. Their auditory cortex is organized into core, belt and parabelt regions as in humans (Belin et al., 2018), which feed higher-order temporal areas involved in complex sounds and conspecific vocalizations processing (Kaas & Hackett, 1998, 2000; Petkov et al., 2006, 2008, Joly et al., 2011) and comparative fMRI has mapped homologous vocalization responses across species (Bodin et al., 2021; Joly et al., 212). Moreover, oxytocin also modulates social attention (Chang & Platt, 2014) and facial-expression responses in macaques (Liu et al., 2015), which makes the macaque a powerful model for testing how OT modulates primate sensory processing.

Here, we tested whether systemic OT modulates auditory processing in a context-dependent manner in awake rhesus macaques. Using high-resolution fMRI (0.6 mm isotropic voxels) with implanted phased-array receive coils (Janssens et al., 2012), we compared intravenous OT with saline across two tasks. In a unimodal auditory task, conspecific vocalizations (coos, screams and aggressive calls) and non-vocal environmental sounds were presented in isolation. In a separate audiovisual task, the same coos and aggressive calls were presented either alone or alongside lip smack and aggressive faces, in congruent and incongruent pairings. Because identical calls occurred in both settings, this design allowed us to address the question whether OT uniformly enhances social auditory responses, or instead modulates sensory responses according to how sounds are encountered.

## Results

Two awake rhesus macaques performed the two tasks across four pharmacological sessions, with saline (CTR) and OT injections counterbalanced (Figure 1). In addition, resting-state scans was acquired after OT and saline injections in the same animals. We first verified that auditory stimulation robustly engaged the expected auditory and auditory-associated cortical network under CTR and OT, then we tested whether OT modulates sound-evoked response amplitude in functionally defined regions of interest (ROIs), whether such change depended on task context and whether it extended to inter-regional coordination.

**Figure 1.**
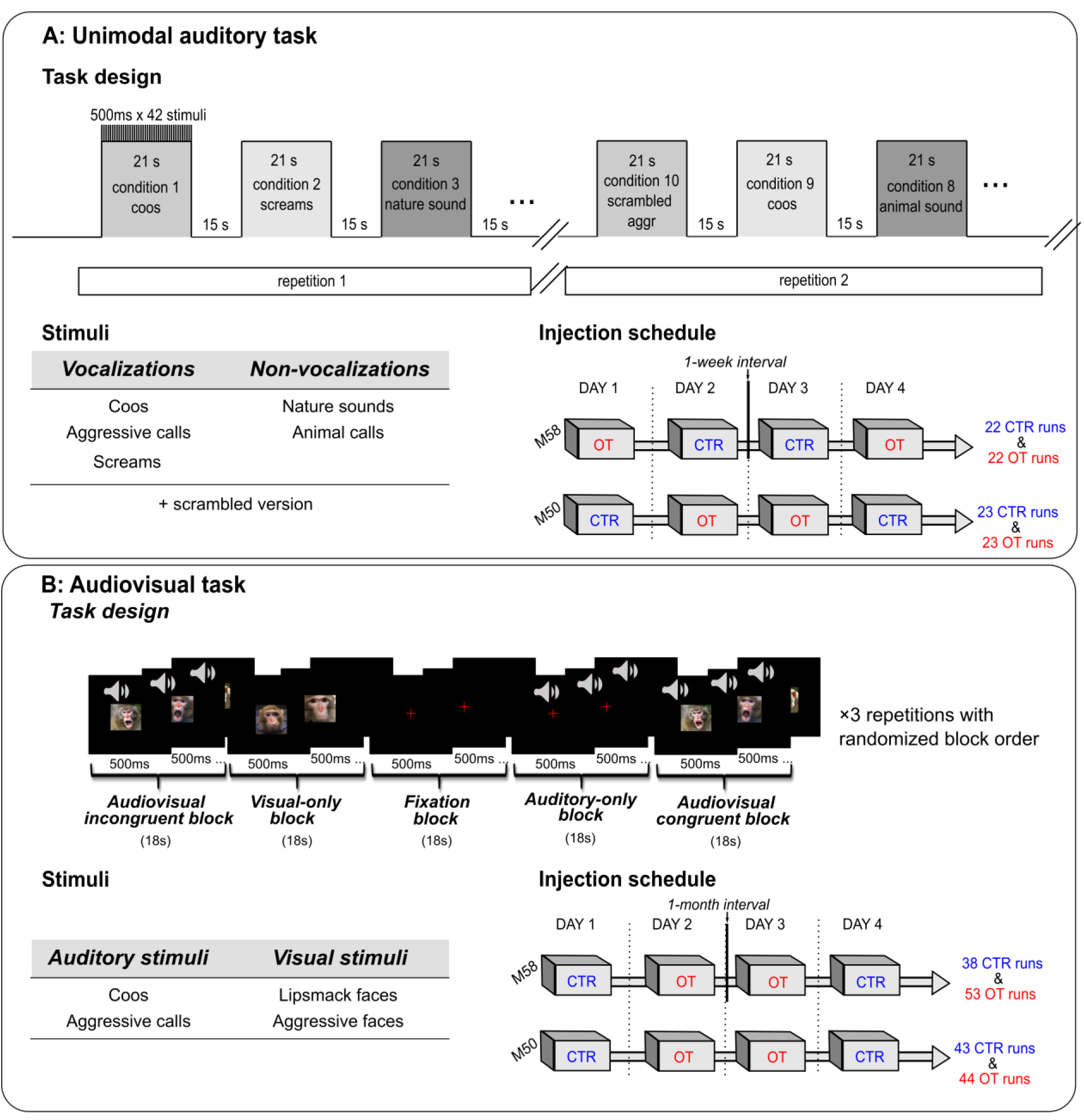
Experimental design and injection schedule for the unimodal auditory and audiovisual tasks. (A) Unimodal Auditory task. During a fixation-task, monkeys listened passively to blocks of vocalizations (coos, aggressive calls and screams) and non-vocal sounds (animal and nature sounds), as well as scrambled versions of each category. Each 21-s sound block contained 42 stimuli of 500 ms and was followed by 15 s of silence. Each condition was presented twice per run, yielding 20 sound blocks per run. Saline (control) and OT injections were administered intravenously across four recording days using a counterbalanced schedule, yielding 22 CTR and 22 OT runs for M58 and 23 CTR and 23 OT runs for M50. (B) Audiovisual task. Blocks of visual-only, auditory-only, audiovisual congruent and audiovisual incongruent social stimuli were presented, with fixation blocks as baseline. Auditory stimuli consisted of coos and aggressive calls, while visual stimuli consisted of lipsmack and aggressive faces. Each block lasted 18 s and the block sequence was repeated three times per run with randomized block order. Control and oxytocin injections were administered across four recording days, yielding 38 CTR and 53 OT runs for M58 and 43 CTR and 44 OT runs for M50. Note that the coos and aggressive calls used in the two tasks were identical. The unimodal auditory task was adapted from Froesel et al., 2025 and the audiovisual task derived from Froesel et al., 2022, 2024.

### Oxytocin attenuates sound-evoked responses in the unimodal auditory task

In the unimodal auditory task, all sound blocks were first contrasted with silence in a combined-monkey analysis. Both CTR and OT sessions elicited widespread activation along the lateral sulcus, covering auditory core and belt regions, superior temporal and insular territories, as well as somato-parietal, premotor and prefrontal regions (Figure 2; *p* < 0.05, family-wise error (FWE) corrected). Auditory stimulation thus engaged a similar distributed network under both injections.

**Figure 2.**
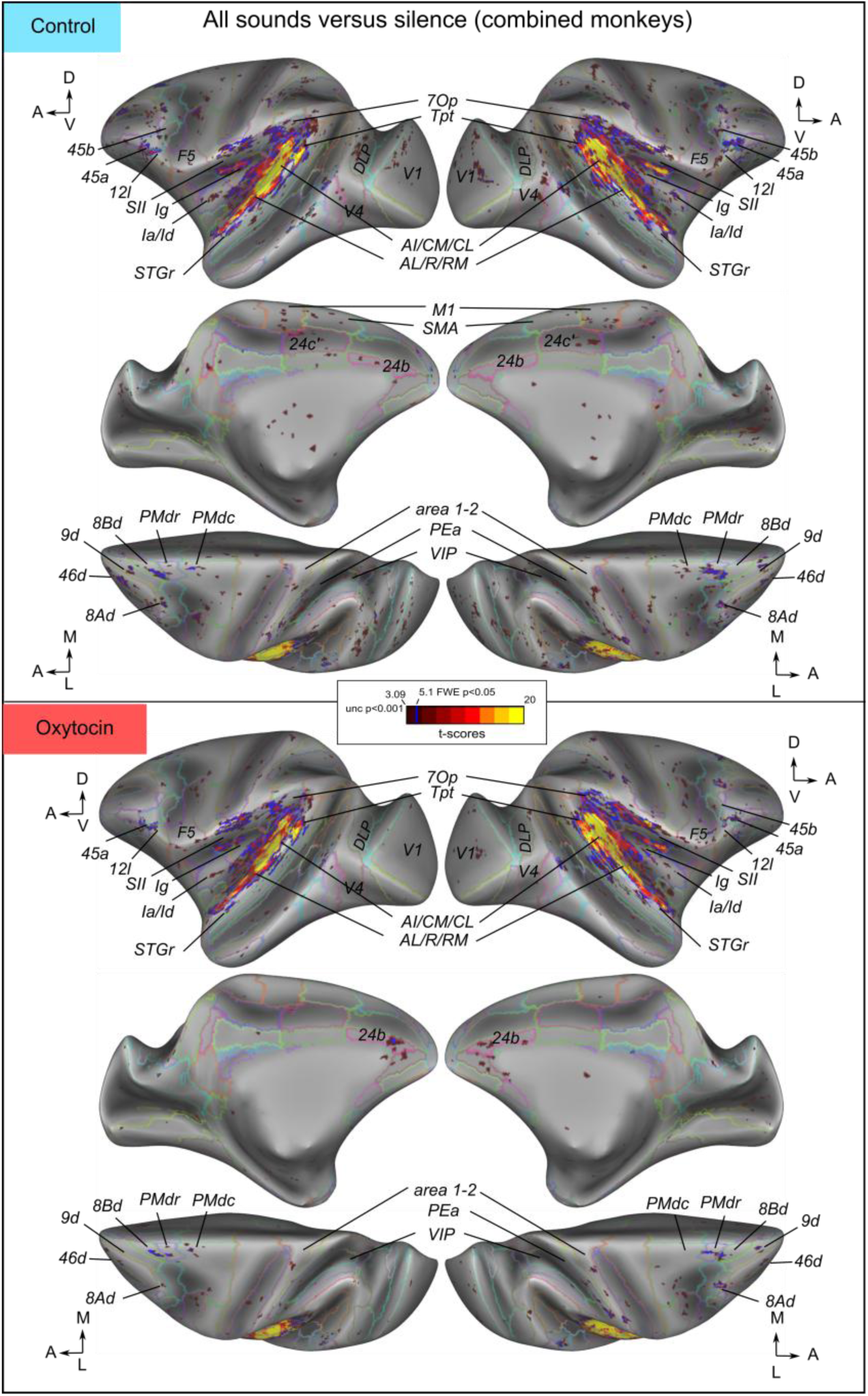
Sound-evoked cortical activation under CTR and OT injections during the unimodal auditory task. Surface projections show SPM *t*-score maps for the contrast all sounds versus silence, displayed separately for CTR/saline and OT sessions in a combined-monkey analysis. Maps are projected onto inflated left and right MEBRAINS template surfaces (Balan et al., 2024). Warm colors indicate sound-evoked activation relative to silence. Statistical maps are displayed at a voxel-wise threshold of *p* < 0.001 uncorrected (*t* > 3.09), with a minimum cluster extent of *k* ≥ 5 contiguous voxels. Blue outlines indicate clusters surviving *p* < 0.05 family-wise error (FWE) correction (*t* > 5.1).

To quantify the OT effect within anatomically interpretable regions, we performed a ROI-based analysis. ROIs were defined from the auditory activation contrast collapsed across injections and subjects and constrained by the RetCHARM atlas (Fitzgerald et al., 2026), so that their selection did not depend on the pharmacological comparison. This yielded 71 hemisphere-specific auditory-responsive cortical ROIs, of which 31 had a counterpart in both hemispheres. These 31 bilateral ROIs were used for the primary ROI analysis and are displayed in Figure 3, where we averaged the homotopic left and right ROIs for display only. Percent signal change (PSC) was extracted from each ROI and run and analyzed using factorial ANOVAs including factors Injection type, Sound category, Hemisphere and Monkey, with FDR correction across the 31 ROIs.

**Figure 3.**
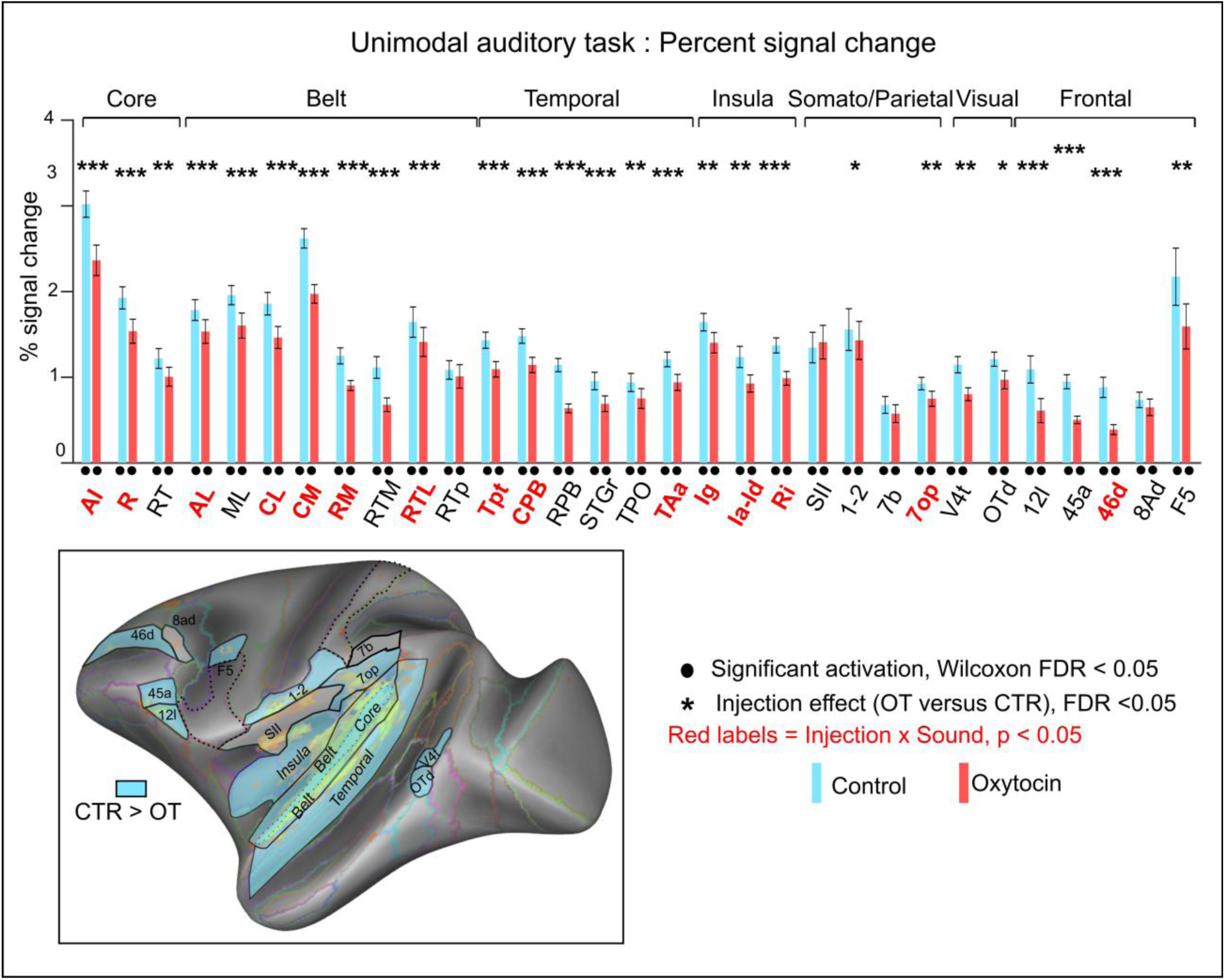
Bilateral auditory-responsive ROI analysis during the unimodal auditory task (pooled analysis). Percent signal change (PSC) in functionally defined auditory-responsive cortical ROIs during the unimodal auditory task, shown for the pooled analysis across both monkeys. Bars represent the mean PSC across the five intact sound categories (aggressive calls, coos, screams, animal sounds, and natural sounds) relative to silence, for control (CTR, blue) and oxytocin (OT, red) sessions. For each ROI, values were averaged across homotopic left and right ROIs. Error bars denote SEM across runs. ROIs are organized from early auditory regions (Core and Belt) to higher-order associative regions (temporal, insular, somato-parietal, visual, and frontal areas). Black dots below individual bars indicate responses significantly different from zero within each condition (Wilcoxon signed-rank tests, FDR-corrected at *p* < 0.05). Asterisks above paired bars indicate a significant main effect of Injection (ANOVA, FDR-corrected across ROIs). Red bold ROI labels indicate regions showing an Injection × Sound interaction (uncorrected *p* < 0.05). The inset provides a schematic cortical summary of the bar plot results, highlighting the ROIs showing stronger responses in CTR than OT sessions.

In both sessions, sounds elicited significant responses relative to baseline in most bilateral ROIs. Despite this preserved activation, OT reduced response magnitude across virtually all ROIs (Figure 3; Supplementary Table 1A,B). Specifically, 27 of the 31 ROIs showed a significant main effect of Injection after FDR correction, all in the CTR > OT direction. The attenuation encompassed all auditory core regions (the primary auditory cortex (AI), rostral core (R), and rostrotemporal core (RT)) and all auditory belt regions except the rostrotemporal-polar area (RTp), including the anterolateral (AL), middle lateral (ML), caudolateral (CL), caudomedial (CM), rostromedial (RM), rostrotemporal-medial (RTM), and rostrotemporal-lateral (RTL) belt regions. The effect extended into higher-order temporal cortex, including the temporoparietal area (Tpt), caudal and rostral parabelt (CPB and RPB), rostral superior temporal gyrus (STGr), temporo-parieto-occipital area (TPO), and temporal area TAa, as well as granular, agranular/dysgranular, and retroinsular cortex (Ig, Ia/Id, and Ri). Additional effects were observed in somato-parietal areas 1–2 and 7op, visual association areas V4t and dorsal occipitotemporal cortex (OTd), and frontal areas 12l, 45a, 46d, and F5.

The direction of this effect was highly consistent across monkeys, although its extent differed (Supplementary Figure 1; Supplementary Table 1A). In M50, 26 of 31 bilateral ROIs showed a significant injection effect after FDR correction, compared to 12 in M58, all with lower responses after OT. 10 ROIs reached FDR significance in the pooled analysis and in each monkey separately (AI, CL, CM, RTL, Tpt, RPB, STGr, area 12l, areas 45a and 46d), spanning core and belt cortex, higher-order temporal association cortex, and prefrontal cortex. 25 of the 27 pooled-significant ROIs exhibited the same direction in each animal, indicating that the attenuation was not driven by one animal even though it was stronger and more widespread in M50. Injection × Hemisphere interactions survived FDR correction in 14 of the 31 bilateral ROIs, indicating regional differences in effect size between hemispheres. The hemisphere-specific analysis gave a similar picture, with significant effects in 42 of 71 ROIs in the pooled analysis (41 CTR > OT), 49 of 71 ROIs in M50, and 16 of 71 in M58 (15 CTR > OT) (Supplementary Table 1A,B).

Because MION fMRI depends on cerebral blood volume, we checked whether the OT-induced attenuation could be explained by local changes in signal quality. Temporal signal-to-noise ratio (tSNR) was higher under OT in nearly every ROI, by 7.9% on average in M50 and 12.3% in M58, reflecting both a higher temporal mean and lower temporal variability. The magnitude of the PSC reduction did not covary with the change in tSNR in either monkey (M50, Spearman rho = 0.13, p = 0.48; M58, rho = -0.15, p = 0.43), nor with changes in temporal mean or standard deviation. Thus, neither the magnitude nor the pattern of the signal-quality change can account for the response attenuation.

The size of the reduction varied across sound categories in a subset of regions: 11 of 31 bilateral ROIs showed a significant Injection × Sound interaction after FDR correction (Supplementary Table 1A,B). However, this interaction was less consistent across animals than the main Injection effect and was predominantly driven by M50. We therefore interpret these category-dependent effects cautiously.

We also examined anatomically defined subcortical ROIs from the SARM atlas (Hartig et al., 2021). 17 regions responded to sound in the pooled analysis. Only claustrum showed a significant injection effect after FDR correction, with lower responses under OT bilaterally (Supplementary Table 2). Other effects appeared in individual monkeys but did not replicate across them. Subcortical modulation was thus far more limited than the cortical effect.

### Sound-category information survives the response reduction

To determine whether the reduction in response amplitude was accompanied by altered sound-category information, we performed five-class multivariate decoding of aggressive calls, coos, screams, animal sounds and natural sounds in four bilateral ROI groups (auditory Core, auditory Belt, temporal multisensory and frontal). Sound category was decodable above chance under both injections in all four groups (all FDR-corrected *q* < 0.05; Supplementary Figure 2A,B). Balanced accuracy was lower under OT, and run-level bootstrap intervals for the OT–CTR difference excluded zero in all four groups (Core: ΔBA = −0.091, 95% CI [−0.157, −0.024]; Belt: −0.073 [−0.145, −0.009]; temporal multisensory: −0.136 [−0.205, −0.077]; frontal: −0.112 [−0.192, −0.027]), but session-level exact reassignment tests did not reach significance. Moreover, a classifier trained on one injection condition and tested on the other stayed above chance in all four ROI groups (all FDR-corrected *q* < 0.05; Supplementary Figure 2C), with the highest accuracy in auditory Belt (0.664) and Core (0.626). Thus, sound-category representations survived the response reduction and generalized across pharmacological states.

### Oxytocin effects differ between unimodal auditory and audiovisual contexts

We next asked whether the oxytocin-induced attenuation of cortical fMRI signals observed in the unimodal auditory task was preserved in a richer audiovisual context. We first focused on the auditory-only blocks of the audiovisual task, in which coos and aggressive calls were presented without concurrent visual stimulation, but within a task context that also included visual-only and audiovisual social blocks (Figure 1B). Because the attenuation in the unimodal auditory task was reproduced when the analysis was restricted to coos and aggressive calls, responses could be compared across tasks within the same auditory-defined ROIs.

In stark contrast to the unimodal auditory task, no widespread CTR > OT reduction was observed during the auditory-only condition in the audiovisual task (Figure 4, upper panel; Supplementary Table 1C). Instead, a restricted set of bilateral ROIs responded more strongly under OT. 9 of the 31 ROIs survived FDR correction for the main effect of Injection, 8 in the OT > CTR direction, including auditory core regions AI, R and RT, belt regions AL, ML and CM, insular area Ia/Id, and premotor area F5, whereas area 46d showed the opposite CTR > OT effect (Supplementary Table 1D). Thus, the broad attenuation observed for isolated sounds in the unimodal auditory task was absent when identical vocalizations occurred inside an audiovisual task structure.

**Figure 4.**
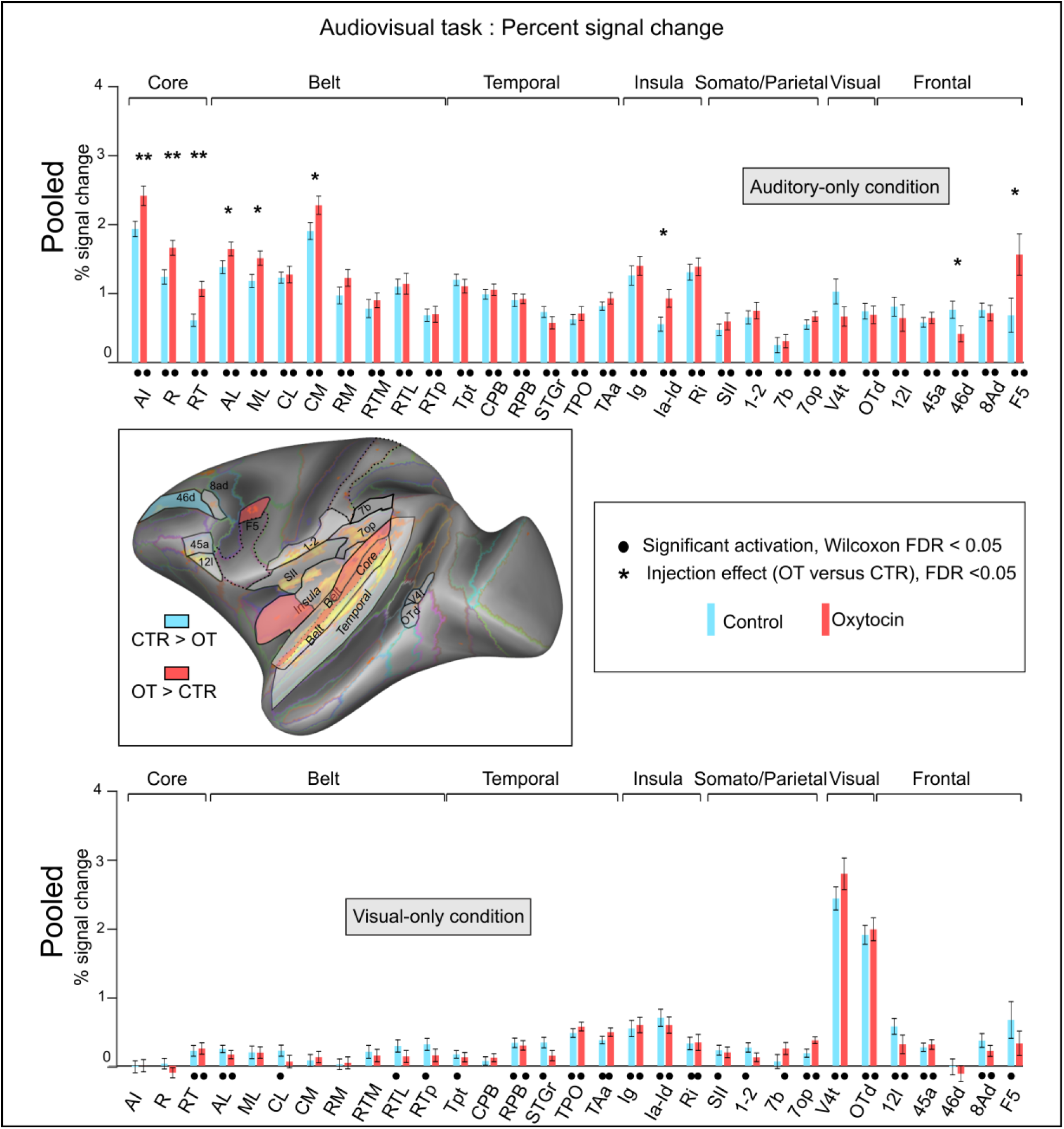
Auditory-defined ROI responses during the audiovisual task. Percent signal change (PSC) was extracted from functionally defined auditory ROIs (defined independently in the unimodal auditory task). Responses are shown separately for the auditory-only condition, in which coos and aggressive calls were presented without visual stimulation, and for the visual-only condition, in which lipsmack and aggressive faces were presented without auditory stimulation. Bars represent mean PSC relative to fixation for control (CTR, blue) and oxytocin (OT, red) sessions. Values were averaged across homologous left and right ROIs. Error bars denote SEM across runs. ROIs are organized from early auditory core and belt regions to higher-order temporal, insular, somato-parietal, visual and frontal regions. Black dots indicate responses significantly different from zero within each condition (Wilcoxon signed-rank tests, FDR-corrected at *p* < 0.05).  Asterisks indicate a significant main effect of Injection after FDR correction across the 31 bilateral ROIs (* *p*FDR < .05; ** pFDR < .01; *** *p*FDR < .001). The cortical inset summarizes ROIs showing a significant main effect of Injection during the auditory-only condition after FDR correction, with cyan indicating CTR > OT and red indicating OT > CTR.

This context-dependent OT > CTR pattern was also present at the individual level, although it differed in spatial extent (Supplementary Table 1D). In M50, 5 bilateral ROIs survived FDR correction, with AI and R showing OT > CTR, and Tpt, V4t and area 46d showing the opposite. In M58, 13 ROIs survived FDR correction, all with larger responses under OT (R, RT, AL, CM, RTL, Tpt, TPO, TAa, Ri, area 7op, area 45a, area 8ad, and F5). The hemisphere-specific analysis yielded a more restricted pattern and is reported in Supplementary Table 1C. Across animals the consistent feature was an increase response concentrated in auditory core and belt regions rather than a widespread shift.

During the visual-only condition, no bilateral auditory-defined ROI showed a significant Injection effect after FDR correction in the pooled analysis (Figure 4 lower panel; Supplementary Table 1E), we therefore did not interpret these effects further. OT effects during visual stimulation were instead assessed directly in independently defined visual-responsive ROIs (Figure 5).

**Figure 5.**
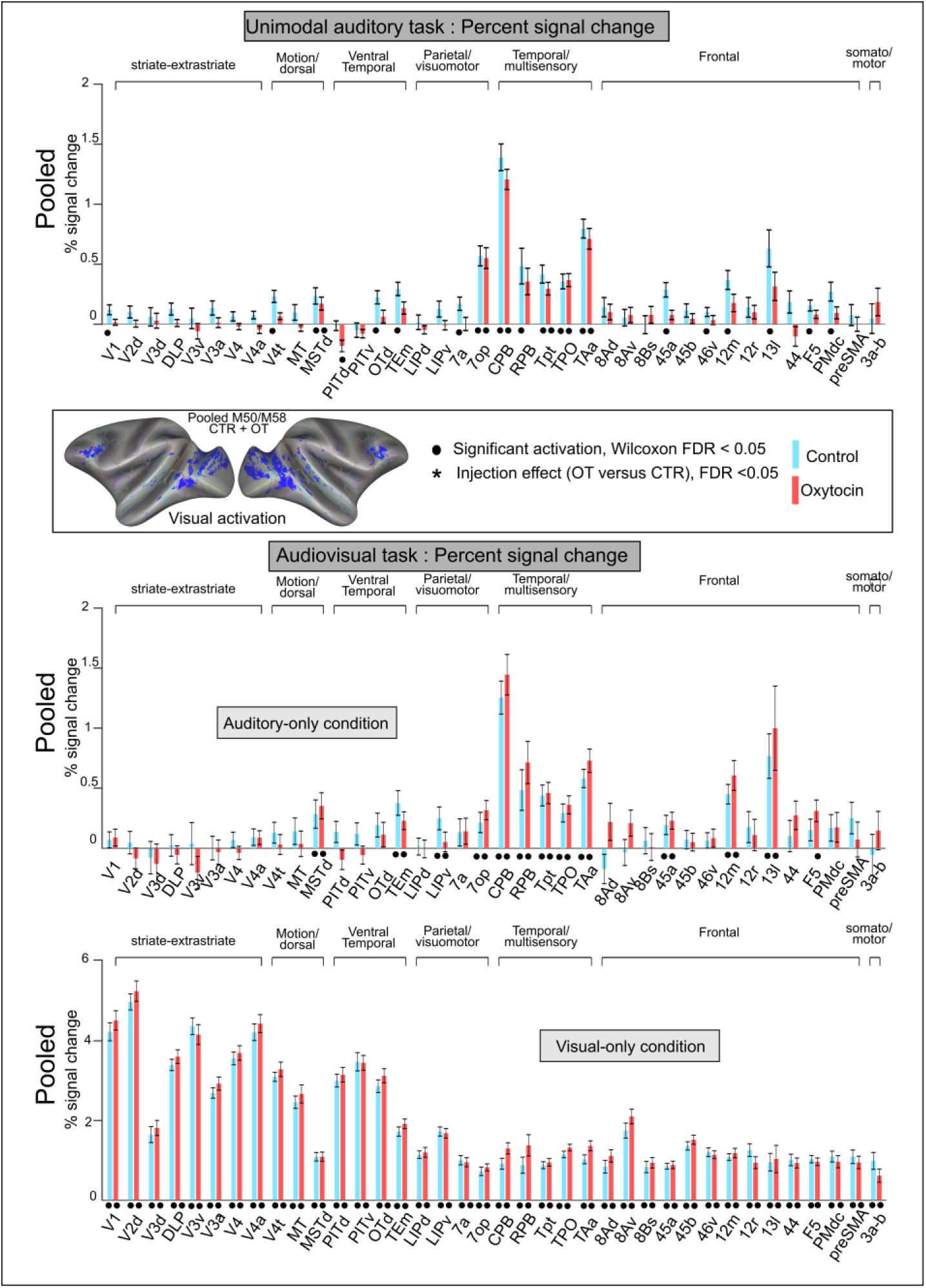
Visual-responsive ROI responses across sensory conditions. Percent signal change (PSC) was extracted from independently defined visual-responsive ROIs during the unimodal auditory task and during the auditory-only and visual-only conditions of the audiovisual task. Bars represent mean PSC relative to fixation for control and oxytocin sessions. Error bars denote SEM. ROIs are organized according to visual and visual-associative cortical subdivisions. Black dots indicate significant responses within condition after FDR correction. Asterisks indicate significant main effects of Injection after FDR correction across ROIs. Visual-responsive ROIs were independently defined from visual-only stimulation relative to fixation in the audiovisual task (CTR and OT sessions and both monkeys pooled; FWE-corrected *p* < .001) and anatomically constrained using RetCHARM. Red bold ROI labels indicate regions exhibiting an Injection × Hemisphere interaction.

Signal-quality analyses again provided no explanation for the observed OT effects during the audiovisual task. OT–CTR differences in PSC during the auditory-only condition did not covary with tSNR changes in either monkey (M50, Spearman rho = −0.11, p = 0.56; M58, rho = 0.15, p = 0.43). Moreover, the auditory-only and visual-only conditions shared the same run-level tSNR estimates while producing different functional effects.

### Oxytocin leaves visual cortex unchanged

To test whether the context-dependent OT effect reflected a general modulation of sensory cortex, as a global vascular or arousal-related influence might produce, we examined visual-responsive ROIs defined from visual-only blocks relative to fixation across both Injection conditions and monkeys and constrained by RetCHARM. These ROIs responded robustly under both injections during visual stimulation (Supplementary Figure 3), but showed no significant injection effect after FDR correction in any condition (Figure 5). The same held when these ROIs were probed during the unimodal auditory task and during auditory-only blocks in the audiovisual task, where they were not engaged.

### Multivoxel representations of task conditions are unchanged

During the audiovisual task, we performed three-class decoding of auditory-only (A), audiovisual-congruent (VAC) and audiovisual-incongruent (VAI) conditions within the same auditory Core and Belt, temporal multisensory and frontal ROI groups. Three-condition accuracy exceeded the 33.3% chance level under both CTR and OT conditions in all four ROI groups (all permutation P=0.001, q=0.001; Supplementary Figure 4 and Supplementary Table 5). Performance was highest in temporal multisensory regions and nearly identical across injections (CTR: 0.639; OT: 0.641). Recall of the auditory-only condition also stayed above chance for both conditions everywhere (all p = 0.001, q = 0.001), reaching 0.854 under CTR and 0.872 under OT in temporal multisensory regions. Exact session-level reassignment tests showed no Injection effect on overall three-condition balanced accuracy or auditory-only recall (all exact p ≥ 0.4, p ≥ 0.6). Thus, auditory-only blocks remained distinguishable from both audiovisual conditions under both CTR and OT, without a detectable change in the multivoxel code.

### Oxytocin reconfigures intrinsic and evoked inter-regional coordination

To examine whether the modulation of task-evoked responses was also expressed in intrinsic network reconfiguration, we analyzed resting-state functional connectivity across ROI groups defined a priori from RetCHARM and SARM atlas regions according to established anatomical and functional subdivisions (Figure 6A; Supplementary Figure 5A; Supplementary Table 6). OT strengthened the pre-existing positive coupling between auditory Core and Belt regions and between the auditory network and visual-motion areas. It attenuated negative couplings with early and extended visual networks and with dorsal prefrontal cortex towards zero. OT also increased near-zero connectivity between auditory cortex and ventral lateral prefrontal cortex (VLPFC) to a significant positive level, weakened negative visual coupling with medial prefrontal cortex and the amygdala, reduced positive auditory coupling with the claustrum/periaqueductal gray (PAG) group and with dorsal premotor cortex, and reduced insula–amygdala coupling. All 11 pooled OT–CTR effects described above survived FDR correction and showed the same direction in at least three of the four monkey-by-hemisphere combinations, with five effects concordant in all four combinations, and each survived FDR correction in at least one monkey when analyzed separately (9 in M50 and 2 in M58). Because each injection condition was represented by a single resting-state session per monkey, these analyses do not provide independent session-level replication of the pharmacological manipulation. Nevertheless, the directional consistency across monkeys and hemispheres, and the satisfactory within-session reproducibility of run-wise connectivity patterns (0.578 to 0.737 across connection families) support a reliable OT-associated reconfiguration of large-scale functional connectivity within the observed sessions. Full statistics are provided in Supplementary Table 6.

**Figure 6.**
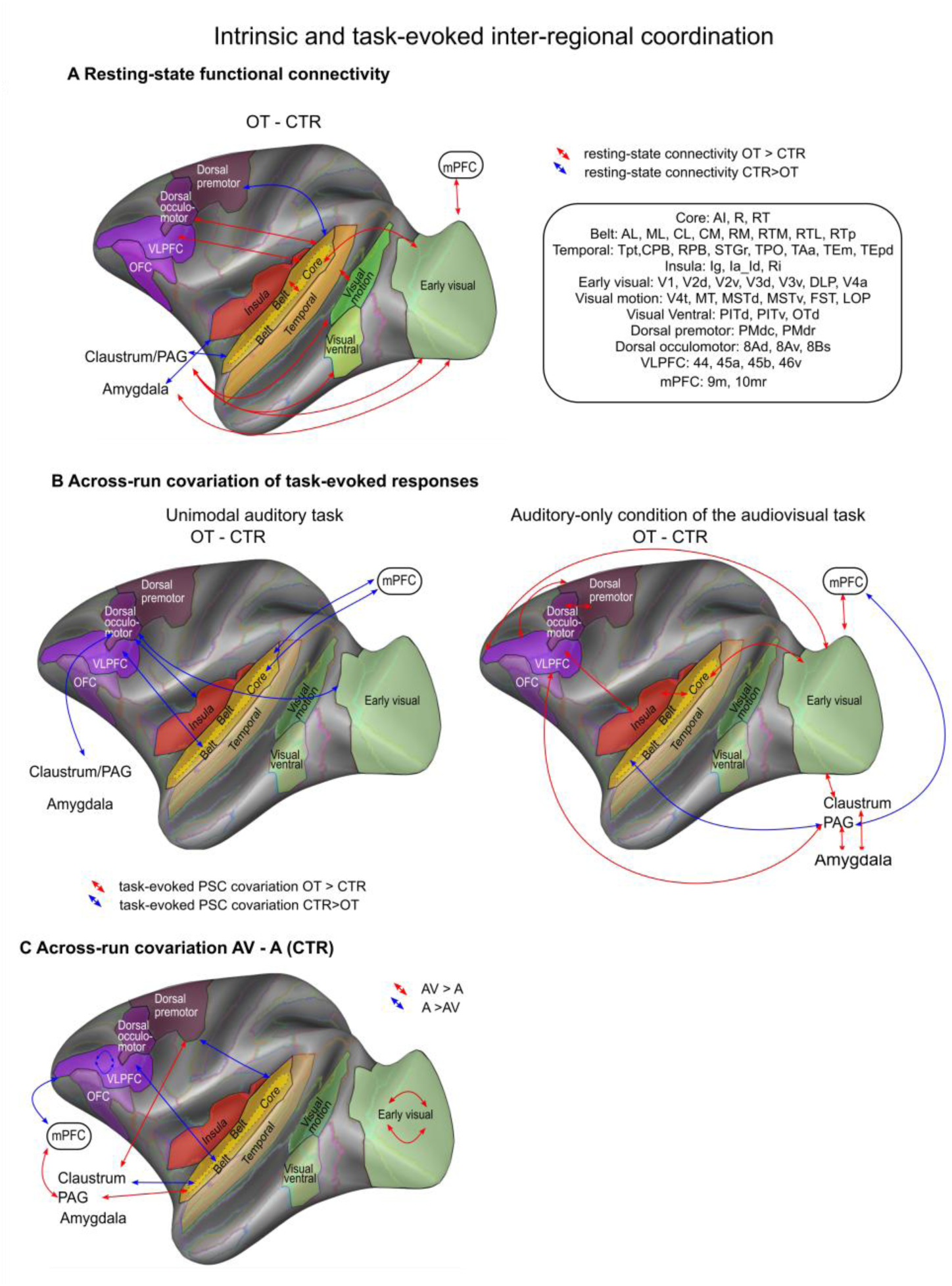
Intrinsic and task-evoked inter-regional coordination. **A)** Resting-state functional connectivity differences between oxytocin (OT) and control (CTR) across predefined ROI groups. Red and blue arrows indicate significantly stronger connectivity under OT and CTR, respectively, after FDR correction across the predefined ROI-group connections. **B)** Across-run covariation of task-evoked percent signal change (PSC) during the unimodal auditory task and the auditory-only condition of the audiovisual task. Brain schematics summarize ROI-group pairs containing ROI-pair Injection effects surviving BH-FDR correction within the corresponding ROI-group pair. Red and blue arrows indicate greater covariation under OT and CTR, respectively. **C)** Direct comparison of across-run PSC covariation between the auditory-only condition of the audiovisual task (AV) and the unimodal auditory task (A) under control (saline) conditions. Red and blue arrows indicate ROI-group pairs containing ROI-pair Task effects surviving BH-FDR correction, with red denoting greater covariation in AV than A and blue denoting greater covariation in A than AV. Circular arrows denote significant within-group effects. Across-run covariation reflects covariation of task-evoked response amplitudes across runs and should not be interpreted as temporal functional connectivity. Only FDR-supported effects are shown; the corresponding ROI-group and ROI-level matrices are provided in the Supplementary Information.

We next asked whether this reconfiguration extended to the coordination of task-evoked response amplitudes by examining across-run covariation of PSC between predefined ROIs, a measure distinct from temporal functional connectivity, tested with run-level mixed-effects models (Figure 6B; Supplementary Figure 5B and 7; Supplementary Table 7). During the unimodal auditory task, 10 ROI pairs survived FDR correction, all showing lower covariation under OT than CTR and concentrated in auditory–prefrontal relationships, including RT–area 45a, RT–area 46v, AI–area 9m, R–area 9m, CM–area 9m, RM–area 9m and CL–area 10mr, with additional effects involving V3d–area 8Av, Ia/Id–area 8Bs and area 8Bs–claustrum. In contrast, during the auditory-only condition of the audiovisual task, 18 ROI pairs survived FDR correction, of which 16 showed greater covariation under OT across auditory–visual, auditory–insular, visual–VLPFC, early visual–claustral, insular–VLPFC, prefrontal–premotor and subcortical relationships. Directly comparing the two tasks under control conditions revealed 14 ROI-pair differences surviving FDR correction (Figure 6C; Supplementary Figure 5C), with the clearest structured effects consisting of increased covariation within the early visual network and reduced covariation between auditory Core and the claustrum. Thus, OT induced both response magnitude and inter-regional coordination changes depending on sensory context.

## Discussion

In this study, we investigated whether systemic oxytocin modulates auditory processing in macaques as a function of sensory context. We found that systemic oxytocin produced opposite effects on the same vocalization stimuli depending on how they were presented. In the unimodal auditory task, isolated sounds were broadly attenuated across auditory core, belt and higher-order associative regions, whereas in an audiovisual task, these calls escaped attenuation and instead OT evoked increased responses in discrete auditory core and belt regions when they were embedded in a structured, cross-modal setting. Stimulus identity cannot explain this reversal, as the attenuation persisted when the unimodal analysis was restricted to the coos and aggressive calls shared across tasks. The context dependence also extended to interregional coordination, which showed markedly different OT-related changes across the two tasks. Together, these results suggest that oxytocin gates sensory responses and their network-level coordination in a context-dependent manner rather than uniformly enhancing or suppressing sensory processing.

The direction of this modulation is consistent with growing evidence that OT effects depend on contextual and motivational factors rather than uniformly enhancing social cues (Bartz et al., 2011; Shamay-Tsoory & Abu-Akel, 2016). In the unimodal task, sounds were presented in isolation during passive fixation and were not embedded in a predictive audiovisual interaction thus carried no explicit behavioral relevance. Under this condition, OT produced a broad attenuation of auditory response magnitude. Importantly, this attenuation was not restricted to a particular class of stimuli: category-dependent effects were relatively limited and primarily reflected differences in the magnitude, rather than the direction, of the OT effect. Thus, the dominant pattern was a general reduction in responses to isolated auditory stimuli rather than a selective suppression of specific vocalization categories. Recent rodent electrophysiology provides a potentially related observation: OT reduces spontaneous activity across auditory, prefrontal and amygdala regions while preserving behaviorally relevant stimulus-evoked information (Jung et al., 2026). Although spontaneous electrophysiological activity and task-evoked fMRI responses are not directly comparable, both findings are compatible with the idea that OT can reduce overall neural responsiveness without eliminating stimulus-specific information. However, because the task was passive and arousal was not measured directly, contributions from changes in orienting or vigilance cannot be excluded.

This broad attenuation interpretation is also consistent with the multivariate results. If OT primarily scaled response magnitude without substantially altering the spatial structure of category representations, category decoding would be expected to remain largely intact despite the lower univariate responses. Consistent with this prediction, the reduction in response magnitude in the unimodal auditory task was accompanied by a decrease in multivariate decoding of call categories, but category information remained decodable above chance under both CTR and OT across auditory Core, Belt, temporal multisensory and frontal networks. Moreover, classifiers trained under one pharmacological condition generalized to the other, particularly in auditory Core and Belt regions, indicating that a substantial component of the sound-category structure was shared across CTR and OT. Similarly, during the audiovisual task, multivoxel patterns discriminated auditory-only, audiovisual congruent and audiovisual incongruent conditions equally well under CTR and OT. We interpret this as evidence that OT reweighted response gain rather than reorganizing categorical representations, consistent with an effect on network-level gain or baseline state rather than on local stimulus selectivity (Jung et al., 2026). This distinction should nevertheless be interpreted cautiously, since preserved fMRI decoding and cross-treatment generalization do not establish that perceptual performance or the underlying neuronal code was unchanged.

Subcortical effects were more restricted, with the bilateral claustrum showing the clearest OT-related attenuation. The claustrum has been implicated in auditory and multisensory processing (Froesel et al., 2024), resilience to sensory distraction (Atlan et al., 2018), and more broadly, in cognitive control through the flexible coordination of cortical networks (Madden et al., 2022). These functions make the claustrum particularly relevant to context-dependent sensory selection, although the broader subcortical pattern was heterogeneous and should be interpreted cautiously. More generally, the attenuation was not global: it was strongest across auditory core, belt and auditory-associated regions, was directionally consistent across both monkeys, and was absent in independently defined visual-responsive ROIs during auditory stimulation. Together with the lack of correspondence between OT-related PSC and tSNR changes, this anatomical specificity argues against a simple vascular or global arousal account.

Why did the same calls escape attenuation within an audiovisual context? The critical difference was not concurrent sensory input per se, since the auditory-only condition of the audiovisual task contained no visual stimulus. Rather, the calls were presented within a task structure that also contained visual-only, audiovisual-congruent and audiovisual-incongruent social blocks. Thus, unlike in the unimodal task, auditory cues belonged to a broader social audiovisual environment in which the two streams could be associated by meaning. Previous work with this paradigm shows that visual social context modulates macaque vocalization responses depending on semantic congruence (Froesel et al., 2022) and face–voice integration engages interactions between auditory cortex and the superior temporal sulcus (Ghazanfar et al., 2008). One possibility is that OT increases the salience of stimuli embedded in a socially meaningful multisensory configuration while dampening isolated acoustic events. Alternatively, cross-modal predictability or within-run expectations rather than semantic meaning per se may determine the direction of modulation. Because both tasks were passive and lacked behavioral readouts or physiological measures of arousal we cannot currently distinguish these alternatives. Matched non-social audiovisual pairings, together with physiological measures such as pupil diameter and heart rate, would separate social meaning from multisensory structure.

The network analyses extended this context dependence to interregional coordination. During the unimodal auditory task, OT consistently reduced across-run covariation of task-evoked responses, particularly across auditory–prefrontal relationships, whereas the audiovisual context showed the opposite dominant pattern, with increased covariation across auditory, visual, insular, prefrontal and subcortical systems. Resting-state connectivity provided complementary evidence that OT also reconfigured large-scale network organization in the absence of sensory stimulation, strengthening selected auditory, auditory–visual and auditory–prefrontal relationships while reducing auditory coupling with premotor and subcortical regions. These resting-state effects should nevertheless be interpreted with caution because each injection condition was represented by a single resting-state session per monkey, limiting independent session-level replication of the pharmacological contrast. Recent rodent electrophysiology provides a useful point of comparison. In mice, OT suppressed spontaneous activity across auditory, prefrontal and amygdala regions while preserving stimulus-evoked information and enhancing sensory–mPFC communication (Jung et al., 2026). Although these measures are not directly comparable to our fMRI results, both studies point to OT-dependent changes in sensory–prefrontal coordination rather than a uniform modulation of sensory activity. In macaques, however, the direction of task-evoked effects depended strongly on sensory context. Together, these results indicate that OT modulates not only local response amplitude but also the coordination of distributed networks, with the expression of this modulation depending strongly on sensory context. Importantly, context-dependent differences under CTR were more restricted than the OT-related changes in task-evoked covariation, arguing against a simple amplification of pre-existing task differences.

The receptor architecture of the macaque constrains how such a state could be implemented. Neocortical regions, including auditory cortex, contain little or no oxytocin receptor (OXTR) in macaques, with binding concentrated in subcortical and limbic structures (Freeman et al., 2014). A purely local action of OT in the regions whose responses changed is therefore unlikely. One possibility is that OT acts through subcortical OXTR-enriched nodes whose ascending projections modulate the gain and coordination of cortical processing, with the direction of this modulation depending on the current sensory context rather than on a fixed property of the stimulus.

These findings may help explain why oxytocin-based interventions in autism spectrum disorder (ASD) have produced heterogeneous results. Beyond the core social-communication difficulties, ASD is characterized by substantial variability in sensory reactivity, filtering and multisensory integration (Kawakami & Otsuka, 2021; Marco et al., 2011; Robertson & Baron-Cohen, 2017), and difficulties in integrating local sensory details into broader contextual representations have also been proposed as a core feature of autistic perception (Happé & Frith, 2006). Alterations of the oxytocinergic system, including differences in OT levels and OXTR regulation, have been reported in ASD and have motivated the investigation of OT as a therapeutic target (Gregory et al., 2009; Lefevre & Sirigu, 2016; Lerer et al., 2008; Modahl et al., 1998; Yamasue & Domes, 2018), yet large randomized trials have not supported a uniform benefit (Sikich et al., 2021). Our results suggest one potential source of this heterogeneity. Rather than uniformly enhancing social processing, OT may reweight sensory signals as a function of their embedding in a structured context. OT-based approaches may therefore benefit from paradigms that explicitly manipulate sensory context and behavioral relevance instead of relying only on broad social outcomes.

Several limitations should be considered. The study involved two macaques, one female and one male, precluding population inference and any separation of sex, age or individual factors. M50 showed a more extensive attenuation in the unimodal task, while the audiovisual enhancement was most consistent in M58, and these differences cannot be disentangled from temperament or age. Resting-state CTR and OT data were each acquired in a single session per monkey, limiting independent session-level replication of the pharmacological contrast despite the consistency of several effects across monkeys and hemispheres. In addition, both tasks were passive, with no behavioral readout of perception and no physiological measures such as pupil diameter or heart rate, so interpretations in terms of reduced vigilance or gain remain hypotheses. Moreover, contrast-based percent signal change cannot distinguish reduced evoked responses from an altered baseline state, although the preserved decoding results and the tSNR control argue against a trivial signal-quality explanation.

In conclusion, systemic OT in awake macaques acts as a context-dependent sensory gate. It attenuates auditory responses in a unimodal auditory task while enhancing responses to the same calls when they are embedded in an audiovisual task context. Despite these changes in response magnitude, stimulus information remained decodable and generalized across pharmacological conditions, indicating that OT modulated sensory processing without abolishing categorical representations. Changes in interregional coordination provided convergent evidence that this modulation extends beyond local response amplitude to distributed network coordination. Together, these findings show that OT does not impose a fixed gain change on sensory processing but modulates sensory responses and their coordination as a function of context. This context dependence may be relevant to the heterogeneous effects of OT reported in humans, particularly where sensory filtering and multisensory integration vary across individuals.

## Materials and Methods

### Subjects

Two adult rhesus monkeys (Macaca mulatta), one female (M50; 6kg; 13 years old) and one male (M58; 9.6kg; 7 years old) participated in the experiment.

Ethics: These animals were raised in captivity, experiencing social interactions primarily with their own species in group housing and with humans during experiments and maintenance of the facilities. All animals were held at the primate facility of the KU Leuven Medical School. The animals were group-housed in cages sized 16 – 32 m^3^, which encourages social interactions and locomotor behavior. The environment was enriched by foraging devices and toys. The animals were fed daily with standard primate chow supplemented with fruits, vegetables, bread, peanuts, cashew nuts, raisins and dry apricots. The animals were water restricted but could work every day until satiated. During non-working days, they received water in their living quarters. Animal care and experimental procedures met the National Institute of Health’s Guide for the Care and Use of Laboratory Animal guidelines, the European legislation (Directive 2010/63/EU) and were approved by the Animal Ethics Committee of KU Leuven.

### Experimental design and stimuli

Monkeys performed two passive fixation tasks during pharmacological fMRI: an unimodal auditory task and an audiovisual task (Figure 1). In both tasks, animals were required to maintain fixation on a central fixation cross throughout the run, while auditory and/or visual stimuli were presented in blocks. Eye position was monitored online at 120 Hz using an infrared eye-tracking system (ISCAN). Runs were included in the analysis only if fixation was maintained within a 2 × 2° window centered on the fixation marker for more than 95% of the run duration.

In the unimodal auditory task, sounds were drawn from a stimulus set described in detail previously (Froesel et al., 2025). Briefly, stimuli were composed of ten sound categories: coos, scrambled coos, nature sounds, scrambled nature sounds, screams, scrambled screams, animal sounds, scrambled animal sounds, aggressive calls and scrambled aggressive calls. Each sound block lasted 21 s and consisted of 42 individual 500-ms sounds. Blocks were separated by 15-s silent fixation periods. Each sound category was presented twice per run, yielding 20 sound blocks per run. Block order was counterbalanced across runs using a Latin-square scheme.

In the audiovisual task, auditory and visual social stimuli were presented either alone or in combination (Froesel et al., 2022, 2024). Auditory stimuli consisted of coos and aggressive calls, corresponding to the same sound categories used in the unimodal auditory task. Visual stimuli consisted of lipsmack and aggressive facial expressions, presented centrally around fixation and subtending approximately 4° of visual angle in radius. The task included auditory-only, visual-only, audiovisual congruent and audiovisual incongruent blocks. Congruent audiovisual blocks paired coos with lipsmack faces and aggressive calls with aggressive faces, whereas incongruent audiovisual blocks paired each sound category with the non-matching facial expression. Each block lasted 18 s, and the full block sequence was repeated three times per run with randomized block order.

Visual stimuli were projected onto a translucent screen positioned 57 cm from the monkeys’ eyes using an LCD projector operating at 60 Hz with a resolution of 1400 × 1050 pixels. Auditory stimuli were delivered at approximately 80 dB SPL through MR-compatible headphones custom-adapted for macaques (MR Confon GmbH, Magdeburg, Germany). The headphone system attenuated scanner noise to below 73 dB SPL, following previously described auditory stimulation procedures for macaque fMRI (Baumgart et al., 1998; Joly et al., 2012).

### Oxytocin (OT) Solution Preparation and Injection

Each monkey completed four pharmacological task-fMRI sessions, comprising two saline control sessions and two oxytocin sessions, during which the unimodal auditory and audiovisual tasks were acquired (Figure 1). Injection order was counterbalanced across animals. Resting-state fMRI was acquired in one saline control session and one oxytocin session per monkey. Oxytocin was administered intravenously at a dose of 25.65 IU in 1 mL, 30 min before the first run. Full details of solution preparation and the exact injection schedules are provided in the Supplementary Methods.

### Scanning procedures

All MRI data were acquired on a 3T Siemens PrismaFit scanner. Functional data were collected using implanted phased-array receive coils embedded in each animal’s headpost, positioned directly above the skull to improve sensitivity for submillimeter fMRI measurements. A 16-channel receive coil was used for M58 and a 10-channel receive coil for M50 (Janssens et al., 2012). Functional scans were acquired with a custom-built local single-loop transmit coil. To increase functional contrast and signal-to-noise ratio, monkeys received an intravenous injection of monocrystalline iron oxide nanoparticles, Molday ION, before each functional scanning session. The contrast agent was administered through the saphenous vein at a dose of 9–11 mg/kg while animals were awake and seated in the scanner chair, following established MION-enhanced fMRI procedures in macaques (Leite et al., 2002; Vanduffel et al., 2001).

#### Anatomy

High-resolution anatomical images were acquired before the functional scanning sessions. Monkeys were anesthetized with ketamine–xylazine for M50 and ketamine–medetomidine for M58. Anatomical scans were acquired using the scanner body transmit coil and a custom-built local single-loop receive coil. For each animal, multiple T1-weighted 3D images were acquired at 0.4 mm isotropic resolution using an MPRAGE sequence. Thirteen T1-weighted images were acquired for M50 and eleven for M58. Sequence parameters were: TR = 2700 ms, TE = 3.5 ms, inversion time = 882 ms, flip angle = 9°, field of view = 104 × 128 mm, matrix size = 250 × 320, and 208 slices. Additional T2-weighted 3D images were acquired to improve tissue segmentation and to assist the identification of dura and blood vessels during surface reconstruction. Five T2-weighted images were acquired for M50 and four for M58, using the same field of view and matrix size as the T1-weighted images. T2-weighted images were acquired with a SPACE sequence: TR = 3200 ms, TE = 456 ms, turbo factor = 131, echo spacing = 6 ms, and no fat suppression. Anatomical data were processed using an adapted non-human primate version of the Human Connectome Project pipeline (Autio et al., 2020), following procedures developed for high-resolution macaque anatomical reconstruction.

#### fMRI

Functional images were acquired in awake macaques during the unimodal auditory and audiovisual tasks using MION-enhanced gradient-echo echo-planar imaging. In both tasks, functional images were acquired at 0.6 mm isotropic resolution and covered the whole brain. Acquisition parameters were identical across tasks: TR = 3000 ms, TE = 21 ms, multiband factor = 2, in-plane acceleration factor = 3, field of view = 84 × 84 mm, matrix size = 140 × 140, and flip angle = 90°. The number of slices was adjusted to each animal’s brain size, resulting in 90 slices for M58 and 80 slices for M50. Each run started with dummy volumes to allow signal stabilization; these volumes were discarded before statistical analysis.

For the unimodal auditory task, each run comprised 243 acquired volumes after the dummy scans. In total, 44 runs were acquired in M58, including 22 control and 22 oxytocin runs, and 46 runs were acquired in M50, including 23 control and 23 oxytocin runs. Only runs with fixation performance above 95% within the 2 × 2° fixation window were included in the analysis.

For the audiovisual task, each run comprised 115 acquired volumes from which we removed the dummy scans. In total, 91 runs were acquired in M58, including 38 control and 53 oxytocin runs, and 87 runs were acquired in M50, including 43 control and 44 oxytocin runs. As for the unimodal auditory task, only runs with fixation performance above 90% within the 2 × 2° fixation window were included in the analysis, and retained runs had fixation performance above 95%.

Resting-state fMRI was acquired using MION-enhanced gradient-echo echo-planar imaging. For M50, images were acquired with TR = 2400 ms, TE = 21 ms, multiband factor = 2, in-plane acceleration factor = 3, a 140 × 140 × 80 matrix and a flip angle of 85°. Each run comprised 275 volumes, and 14 runs were acquired under each injection condition. For M58, images were acquired with TR = 1500 ms, TE = 21 ms, multiband factor = 2, in-plane acceleration factor = 3, a 140 × 140 × 78 matrix and a flip angle of 75°. Each run comprised 600 volumes, and 10 runs were acquired under each injection condition. For resting-state acquisitions, each injection condition was represented by one acquisition session per monkey.

### Preprocessing

Functional data from the task and resting-state acquisitions were preprocessed using a combination of FSL, SPM12, the JIP analysis toolkit, Advanced Normalization Tools (ANTs) and Connectome Workbench (Jenkinson et al., 2012). Preprocessing was performed separately for each monkey and each task before normalization to a common template space.

For each subject, a reference EPI template was generated from the first ten volumes acquired on the first scanning day. EPI volumes were skull-stripped and corrected for magnetic field inhomogeneities using fieldmap-based distortion correction. Head motion was corrected using rigid-body registration with six degrees of freedom. To further reduce residual frame-to-frame geometric distortions, we applied slice-wise nonlinear distortion correction using a B-spline grid registration approach, with all volumes aligned to the individual EPI reference template (Kolster et al., 2009). Functional images were then spatially smoothed with a Gaussian kernel of 1 mm full width at half maximum.

Task-based functional time series from the unimodal auditory-only and audiovisual tasks were denoised using GLMdenoise (Kay et al., 2013). For each run, time series were first converted to percent signal change. GLMdenoise identifies a pool of noise voxels with little or no task-related variance and extracts nuisance components from these time series using principal component analysis. These components were then progressively added to the task GLM, and the optimal number of nuisance regressors was selected based on cross-validated improvement in model fit. The selected nuisance components were regressed from the functional data to obtain denoised task time series.

Resting-state runs underwent the same geometric correction, coregistration and normalization procedures. Temporal filtering and nuisance-signal correction using white-matter and cerebrospinal-fluid masks were performed separately for each run.

The preprocessed functional data were coregistered to each animal’s high-resolution anatomical image using a combination of linear and nonlinear registration with ANTs. Anatomical images were processed using an adapted non-human primate version of the Human Connectome Project pipeline (Autio et al., 2020; Glasser et al., 2013). Individual anatomical and functional data were then normalized to the MEBRAINS template space (Balan et al., 2024), allowing combined analyses across the two monkeys

### First-Level GLM Analysis with SPM

For each individual fMRI run, a General Linear Model (GLM) was specified and estimated using SPM (Statistical Parametric Mapping). Fixed effect analyses were performed for each run of each monkey using head motion as regressor of no interest and a MION hemodynamic response function to convolve the signal (Vanduffel et al., 2001).

For the unimodal auditory-only task, first-level contrast images were generated for each sound category relative to silence, for all sounds relative to silence, for vocalizations relative to silence, and for non-vocalizations versus silence.

For the audiovisual task, first-level GLMs included separate regressors for auditory-only, visual-only, audiovisual congruent and audiovisual incongruent blocks, with additional regressors modeling stimulus category where appropriate. First-level contrasts were generated for each task condition relative to fixation, as well as for contrasts comparing auditory-only, visual-only and audiovisual conditions.

### Temporal Signal-to-Noise Ratio (tSNR) Analysis

To assess whether injection-related functional differences could be explained by differences in signal quality, we performed a voxel-wise temporal signal-to-noise ratio (tSNR) analysis as a quality-control analysis. For each functional run, tSNR was computed as the temporal mean of the preprocessed and denoised MION time series divided by its temporal standard deviation, using FSL fslmaths. This yielded one tSNR map per run. Run-level tSNR maps were compared between control and oxytocin sessions using voxel-wise two-sample *t*-tests implemented in AFNI 3dttest++. Statistical maps were corrected for multiple comparisons using false discovery rate correction at *q* < 0.05. To quantify the magnitude of injection-related tSNR differences, voxel-wise Cohen’s *d* effect-size maps were computed using the pooled standard deviation of the two injection conditions. In addition, the temporal mean signal and temporal standard deviation used to compute tSNR were retained separately to determine which component contributed to injection-related changes in tSNR. For each monkey and task, maps of tSNR, temporal mean signal, and temporal standard deviation were averaged separately across CTR and OT runs, and mean values were extracted from the same functionally defined cortical ROIs used for the PSC analyses. Analyses were performed in both the auditory-responsive ROI set defined from the auditory-only task and the visual-responsive ROI set defined from visual stimulation in the audiovisual task. For the unimodal auditory task, functional effects were summarized as the OT–CTR PSC difference averaged across the five intact sound categories. For the audiovisual task, OT–CTR PSC differences were calculated separately for the auditory-only and visual-only conditions. Because these conditions occurred within the same audiovisual runs, tSNR, temporal mean signal, and temporal standard deviation were calculated at the run level and were therefore common to the auditory only and visual only analyses, whereas the corresponding PSC difference was condition-specific. For each monkey, Spearman rank correlations across ROIs were used to assess whether the magnitude of the functional OT–CTR effect covaried with injection-related changes in tSNR, temporal mean signal, or temporal standard deviation. Equal-weight pooled estimates across the two monkeys were additionally calculated descriptively. ROI-wise correlations were interpreted descriptively because anatomical ROIs are not statistically independent spatial observations.

### ROI analysis

Cortical regions of interest (ROIs) were defined using task-evoked functional contrasts independent of the pharmacological comparison, collapsed across Injection conditions and monkeys, and anatomically constrained using the RetCHARM atlas (Fitzgerald et al., 2026). Subcortical ROIs were defined anatomically using the SARM atlas without functional preselection. Percent signal change was extracted using MarsBaR (Brett et al., 2002) and, because MION-related increases in cerebral blood volume decrease the MR signal, was calculated as Fixation − Condition, such that positive values indicated task-evoked activation. Injection and condition effects were analysed at the run level using factorial ANOVAs, whereas activation relative to fixation or silence was assessed using two-sided Wilcoxon signed-rank tests. Benjamini–Hochberg FDR correction was applied within predefined analysis ROI groups. Complete ROI definitions, mask-construction procedures and detailed model specifications are provided in the Supplementary Methods and Supplementary Table 3.

#### Multivoxel pattern analysis of sound-category information

To quantify whether sound-category and task-condition information was preserved, we performed ROI-based multivoxel pattern analysis (MVPA) using linear discriminant analysis with automatic covariance shrinkage. Decoding performance (balanced accuracy and class-specific recall) was evaluated using stratified cross-validation within predefined auditory, temporal, and frontal ROI groups. Significance relative to chance was assessed using nonparametric label permutations, and injection effects (OT vs. CTR) were tested using exact reassignment of complete pharmacological sessions. Full details regarding pattern extraction, cross-validation schemes, and permutation testing are available in the Supplementary Methods.

#### Resting-state functional connectivity analysis

Resting-state functional connectivity was assessed separately in each hemisphere using 83 anatomically defined cortical and subcortical ROIs from the RetCHARM and SARM atlases (Hartig et al., 2021). For each run, pairwise ROI correlations were Fisher-z transformed and summarized within predefined connection ROI groups, yielding one estimate per family, run, monkey, hemisphere and Injection condition. CTR, OT and OT–CTR effects were calculated with equal weighting of the four monkey-by-hemisphere combinations and assessed using stratified permutation tests, with FDR correction across connection ROI groups. Because each Injection condition was represented by one resting-state session per monkey, these analyses were considered exploratory. Full preprocessing, network definitions, reliability, bootstrap and post-hoc node-level procedures are described in the Supplementary Methods.

#### Across-run covariation of task-evoked responses

Across-run covariation of task-evoked PSC was examined across 40 anatomically defined RetCHARM and SARM ROIs grouped into 11 predefined ROI groups. For the unimodal auditory task, the run-level PSC measure was the equally weighted mean response to coos and aggressive calls; for the audiovisual task, the auditory-only PSC was used. For each ipsilateral ROI pair, PSC values were centered within Monkey × Injection × Session × Hemisphere; audiovisual data were additionally centered within sound category because coos and aggressive calls occurred in separate runs. Centered values were rank-transformed and standardized, and Injection effects on covariation were tested using run-level mixed-effects models containing a ROI-response × Injection interaction, with Monkey and Hemisphere as fixed effects and random intercepts for session and run. Descriptive Spearman correlations were calculated separately under CTR and OT, and FDR correction was applied within each predefined connection-family block. Full analysis details are provided in the Supplementary Methods.

## Data and Code Availability

The data and code for the Control unimodal auditory task are available via KU Leuven RDR dataset named High-resolution fMRI that reveals a dorsal brain pathway selective for conspecific vocalizations in macaques (https://doi.org/10.48804/P841JQ). Any additional information required to reanalyze the data reported in this paper is available from the lead contacts upon request.

The data and code for the unimodal auditory task with oxytocin will be also available on KU Leuven RDR as well as the audiovisual task once the data will be fully analyzed and published.

## Acknowledgements

The authors thank C. Fransen, I. Puttemans, C Ulens, A. Hermans, S. Verstraeten, S. Tajmirriyahi, W. Depuydt, and M. De Paep for technical and administrative support. We thank Arthur Lefevre for his advice.

This work received funding from KU Leuven C14/21/111, IDN/20/016, C3/21/027, CELSA/24/020; Fonds Wetenschappelijk Onderzoek-Vlaanderen (FWO-Flanders) 12AF124N, G0E0520N, G0C1920N; and the European Union’s Horizon 2020 Framework Programme for Research H2020, grant EBRAINS2.0., French national research agency (ANR) ANR-20-CE37-0005.

## Author Contributions

Conceptualization, M.F. Q.Z. and W.V.; Stimuli preparation, M.F; Methodology, M.F., H.W., W.V., and Q.Z; Data acquisition, M.F., and Q.Z; Analyses, M.F. ; Funding, W.V., Q.Z. and M.F.; Writing, M.F, W.V. and Q.Z.; Review and editing, Q.Z., M.F., H.W. and W.V.; Supervision, W.V. and Q.Z.

## Competing Interests

No competing interest

## Notes

### Competing Interest Statement

The authors have declared no competing interest.

