## Supplemental Data for "Sensory context gates oxytocin effects on auditory processing in macaques"

### Supplementary data

#### Supplementary methods

##### Oxytocin preparation and pharmacological schedule

A stock solution of oxytocin (OT) was prepared at 4.275 mg/mL by reconstituting 5 mg of OT powder in 1.17 mL of sterile, non-pyrogenic Water for Injection. The stock solution was divided into 15-µL aliquots and stored frozen until use. On each OT experimental day, one aliquot was diluted with 1,485 µL of sterile Water for Injection, yielding a final volume of 1.5 mL. A 1-mL volume of this solution, corresponding to a total dose of 25.65 IU OT, was administered intravenously 30 min before the beginning of the first functional run. During control sessions animals received 1 mL of sterile saline through the same intravenous route at the same pre-run timing. For the task-fMRI experiment, each monkey completed four pharmacological scanning sessions, comprising two saline control (CTR) sessions and two OT sessions. Injection order was counterbalanced across animals, with M50 receiving the sequence CTR–OT–OT–CTR, whereas M58 received the reverse sequence OT–CTR–CTR–OT. Both the unimodal auditory and audiovisual tasks were acquired across these four sessions. Resting-state fMRI was acquired in one saline control session and one OT session per monkey.

##### ROI definition and univariate analyses

###### *Cortical regions of interest*

Cortical regions of interest (ROIs) were defined using functional contrasts independent of the pharmacological comparison. Auditory-responsive ROIs were identified from the *All sounds* > *Silence* contrast in the unimodal auditory task, using data pooled across CTR and OT sessions and across both monkeys. The resulting activation map was thresholded at FWE $p < 0.001$ and intersected with parcels from the RetCHARM macaque atlas to obtain anatomically constrained cortical ROIs. Visual-responsive ROIs were similarly defined from the *Visual only* > *Fixation* contrast in the audiovisual task (thresholded at FWE $p < 0.001$), using data pooled across injections and monkeys, and were constrained using RetCHARM parcels. Subcortical ROIs were anatomically defined from the SARM atlas without functional preselection. The complete auditory cortical, visual cortical and subcortical ROI sets, together with their defining contrasts, are reported in Supplementary Table 3.

###### *Percent-signal-change extraction*

Percent signal change (PSC) was extracted from each run and ROI using MarsBaR. Because increases in cerebral blood volume produce decreases in the MION signal, PSC was calculated as Fixation − Condition, such that positive values represented task-evoked activation. Analyses were performed using both hemisphere-specific estimates and bilateral ROI summaries. For the unimodal auditory task, PSC values were analyzed for the five intact sound categories (aggressive calls, coos, screams, animal sounds and natural environmental sounds). Factorial ANOVAs included Injection, Sound category and their interaction. For the auditory-only condition of the audiovisual task, models included Injection and, where applicable, Hemisphere and Monkey. The same general framework was applied to visual-responsive ROIs during auditory-only and visual stimulation conditions. Pooled analyses combined runs from both monkeys and included Monkey as a fixed factor when relevant. Analyses were also repeated separately in M50 and M58 to evaluate cross-animal consistency. Post hoc control-versus-oxytocin comparisons were performed within individual sound or task conditions when justified by the corresponding main effect or interaction. Activation relative to fixation or silence was assessed using two-sided Wilcoxon signed-rank tests against zero, performed separately for each ROI and injection condition. Multiple comparisons were controlled using Benjamini–Hochberg false-discovery-rate correction applied separately for each ROI set, task, analysis level (pooled or individual monkey), and effect tested. Effects were considered significant at $q < 0.05$.

#### Multivoxel pattern analysis (MVPA)

ROI-based multivoxel pattern analyses were performed in Python using NiBabel and scikit-learn. Analyses were restricted to functional RetCHARM ROIs defined independently of the pharmacological comparison (Supplementary Table 4). Masks included auditory Core regions (AI, R and RT), auditory Belt regions (AL, ML, CL, CM, RM, RTM, RTL and RTp), temporal and multisensory regions (Tpt, CPB, RPB, STGr, TPO and TAa), and frontal regions (45a and 46d). Analyses were performed for individual left and right hemisphere ROIs, bilateral unions of homologous ROIs, and hemisphere-specific and bilateral anatomical-family masks. For each run-by-condition observation, the mean beta value across voxels within the ROI was subtracted from the multivoxel pattern, and the resulting demeaned pattern was normalized to unit Euclidean norm. For each cross-validation fold, the training data comprised all run-by-condition patterns assigned to the training runs, whereas patterns from the remaining runs formed the held-out test data. All condition-specific patterns originating from the same run were kept within the same fold. Voxel-wise means and standard deviations were estimated exclusively from the training observations and subsequently applied to both the training and held-out data. Classification was performed separately for each monkey and Injection condition using linear discriminant analysis with automatic covariance shrinkage. Balanced accuracy was used as the primary measure of decoding performance.

In the unimodal auditory task, multivoxel patterns were extracted from first-level beta images for the five intact sound categories (aggressive calls, coos, screams, animal sounds and natural environmental sounds). Each run contributed five observations, and theoretical chance performance was 20%. Two-fold cross-validation was stratified across the two sessions available for each monkey and Injection condition. Odd-numbered runs from both sessions were used for training and even-numbered runs for testing in the first fold, with the assignment reversed in the second fold.

In the audiovisual task, multivoxel patterns were extracted from the first-level beta images corresponding to the auditory-only (AC), audiovisual-congruent (VAC), and audiovisual-incongruent (VAI) conditions after subtraction of the block-fixation beta image. Each run contributed three observations, and theoretical chance performance was 33.3%. Classification used five-fold run-level cross-validation. In addition to overall three-condition balanced accuracy, class-specific recall was calculated for the auditory-only condition.

Significance relative to chance was assessed using 1000 nonparametric permutations in which condition labels were reassigned within runs while preserving run structure. The complete preprocessing and classification procedure was repeated for every permutation. One-sided permutation p-values were calculated using the plus-one correction, and Benjamini–Hochberg FDR correction was applied across ROIs separately within each task, Injection condition and ROI-analysis level, and separately for monkey-specific and pooled analyses. Confusion matrices were calculated from the combined held-out predictions, and class-specific performance was quantified as recall from the diagonal of the row-normalized confusion matrix. The same permutation procedure was used to assess class-specific recall relative to chance. Descriptive 95% confidence intervals were obtained from 500 run-level bootstrap resamples. For pooled analyses, classification was performed independently in each monkey, and balanced accuracy, recall and the corresponding permutation statistics were averaged with equal weighting across monkeys; voxel patterns were not concatenated across animals. Injection effects were quantified as OT–CTR differences and tested by exact reassignment of complete pharmacological sessions. Within each monkey, two of the four sessions were assigned to pseudo-OT and two to pseudo-CTR, yielding 36 combined assignments across the two monkeys. Two-sided exact p-values were derived from this distribution and FDR-corrected separately for each ROI-analysis level. The same session-level reassignment procedure was applied to Injection effects on class-specific recall. Runs and cross-validation folds were not treated as independent pharmacological replications.

#### Resting-state functional connectivity analysis

##### *Data preprocessing and ROI-to-ROI connectivity*

Resting-state functional connectivity was examined in the two macaques following saline control (CTR) and oxytocin (OT) administration. M50 contributed 14 runs per condition and M58 contributed 10 runs per condition. Data from the left and right hemispheres were analyzed separately. Each injection condition was represented by one acquisition session per monkey. Resting-state runs were processed individually using the geometric correction, motion-correction, coregistration and normalization procedures described for the fMRI data. Nuisance signals were regressed separately from each run. Mean time series were extracted from white-matter and cerebrospinal-fluid masks and combined with the estimated motion regressors. These nuisance regressors were entered into a general linear model using FSL, and the resulting residual four-dimensional images were used for the connectivity analysis. No additional temporal band-pass filtering was applied at this stage.

Cortical and subcortical regions of interest were defined anatomically using the RetCHARM and SARM atlases, respectively. A total of 83 ROIs were included in each hemisphere. For each run, the mean residual time series was extracted from every ROI and standardized to zero mean and unit variance. Pairwise Pearson correlations were then calculated between all ROI time series, yielding one 83×83 functional-connectivity matrix per run. Diagonal values were excluded, and correlation coefficients were transformed using Fisher's r-to-$z$ transformation before statistical analysis.

##### *Network and connection-family definitions*

For the predefined connection-family analyses, ROIs were grouped into anatomically and functionally defined networks of interest. The auditory Core comprised AI, R and RT, whereas the auditory Belt comprised AL, ML, CL, CM, RM, RTM, RTL and RTp. Temporal regions included Tpt, CPB, RPB, STGr, TPO and TAa, and the insular group included Ig, Ia/Id and Ri. Visual regions were divided into early visual, visual-motion and ventral visual networks. An extended visual network was defined as the union of these three visual groups. Frontal regions were divided into orbitofrontal, ventrolateral prefrontal, dorsal/oculomotor prefrontal, medial prefrontal and dorsal premotor networks. Additional subcortical groups included the medial geniculate nucleus, the claustrum/PAG composite and the amygdala. The auditory Core/Belt network was defined as the union of auditory Core and Belt regions. Complete network compositions are reported in the Statistical Source Data. Connectivity was summarized using predefined connection families. For connections between two networks, all Fisher-$z$-transformed edges linking an ROI in the first network to an ROI in the second network were averaged. This procedure produced one connectivity value per family, run, monkey, hemisphere and injection, thereby avoiding the treatment of individual edges from the same correlation matrix as statistically independent observations. Because these measures were based on pairwise correlations, all network relationships were interpreted as undirected functional coupling.

##### *Statistical analyses and quality control*

For each connection family, condition-specific connectivity was first averaged separately within each of the four monkey-by-hemisphere combinations (M50 left, M50 right, M58 left, M58 right). The overall CTR or OT estimate was then calculated as the unweighted mean of these four values, preventing M50 from receiving greater weight because it contributed more runs than M58. Connectivity under CTR and OT was tested separately against zero using two-sided stratified sign-flip permutation tests with 10,000 permutations. At each permutation, the sign of each run-wise score was randomly inverted within its original monkey-by-hemisphere combination, and the equal-weight overall mean was recalculated. Conditional 95% confidence intervals were estimated using 5000 bootstrap samples, with runs resampled separately within each monkey, hemisphere and injection. FDR correction was applied separately across the predefined connection families for CTR and OT.

For direct injection comparisons, the OT–CTR difference was calculated within each of the four monkey-by-hemisphere combinations and then averaged with equal weighting. Statistical significance was evaluated using a two-sided stratified permutation test with 10,000 permutations. Injection labels were shuffled among runs within each monkey-by-hemisphere combination while preserving the original numbers of CTR and OT runs. Conditional 95% confidence intervals were obtained from 5,000 bootstrap samples, with CTR and OT runs resampled separately within each combination. FDR correction was applied across the predefined connection families.

Directional consistency was summarized as the number of the four monkey-by-hemisphere combinations showing the same OT–CTR direction as the overall estimate; it ranged from 0 to 4 and was reported in the form n/4. This measure served as a description of agreement rather than as an additional statistical test.

Within-session reproducibility was evaluated separately for each monkey, hemisphere and injection. For each run, the lower triangle of the 83 × 83 matrix was vectorized, excluding the diagonal, yielding 3,403 unique connections. Leave-one-run-out reliability was calculated as the Pearson correlation between the Fisher-$z$ edge vector of one run and the mean edge vector obtained from all remaining runs from the same monkey, hemisphere and injection. Potentially atypical runs were identified using a robust lower threshold defined as the group median minus $3 \times 1.4826 \times$ the median absolute deviation. This measure was used for quality-control assessment and not as an automatic exclusion criterion. Run-wise connectivity patterns showed satisfactory within-session reproducibility, with median similarities ranging from 0.578 to 0.737. M50 OT run 13 fell below the robust threshold in both hemispheres; because this criterion was used for quality-control assessment rather than automatic exclusion, all runs were retained in the final analyses.

For selected connection families showing significant OT–CTR effects, post-hoc node analyses were used to localize the strongest ROI-level contributions. For each constituent ROI, its contribution was defined as its mean Fisher-$z$ connectivity with all ROIs belonging to the partner network. OT–CTR effects were then assessed using the same equal-weight, stratified permutation and bootstrap framework as the family-level analyses. FDR correction was applied across ROIs within each family and on each side of the corresponding network pair. These analyses localized family-level effects within the same dataset and were not interpreted as independent ROI-level or edge-level confirmation.

Because CTR and OT were each represented by a single resting-state acquisition session per monkey, runs were not considered independent replications of the pharmacological manipulation. The permutation analyses quantified the consistency of connectivity differences across runs, hemispheres and monkeys within the observed sessions. Resting-state findings were therefore interpreted as exploratory evidence of OT-associated network reconfiguration rather than definitive population-level inference or full separation of injection effects from session-specific influences.

#### Across-run covariation of task-evoked responses

Across-run covariation was assessed using the run-level PSC estimates and predefined ROI families described above. All distinct ipsilateral ROI pairs within and between the selected families were tested. For the unimodal auditory task, coo and aggressive-call PSCs were averaged within each run. For the auditory-only condition of the audiovisual task, all runs were retained and sound category was explicitly controlled because coos and aggressive calls occurred in separate runs. PSC values were centered within Monkey × Injection × Session ×

Hemisphere, additionally within sound category for the audiovisual task, then rank-transformed and standardized within Monkey × Injection × Hemisphere. Injection effects on covariation were tested using run-level mixed-effects models containing an ROI-response × Injection interaction, with Monkey and Hemisphere as fixed effects and random intercepts for Session and Run; the audiovisual model additionally included sound category. Descriptive Spearman correlations were calculated separately under CTR and OT within each monkey-by-hemisphere combination and combined with equal weighting after Fisher-$z$ transformation. Δ Fisher $z$ was defined as $z$(OT) − $z$(CTR). FDR correction was applied separately within each predefined connection-family block.

### Supplementary figures

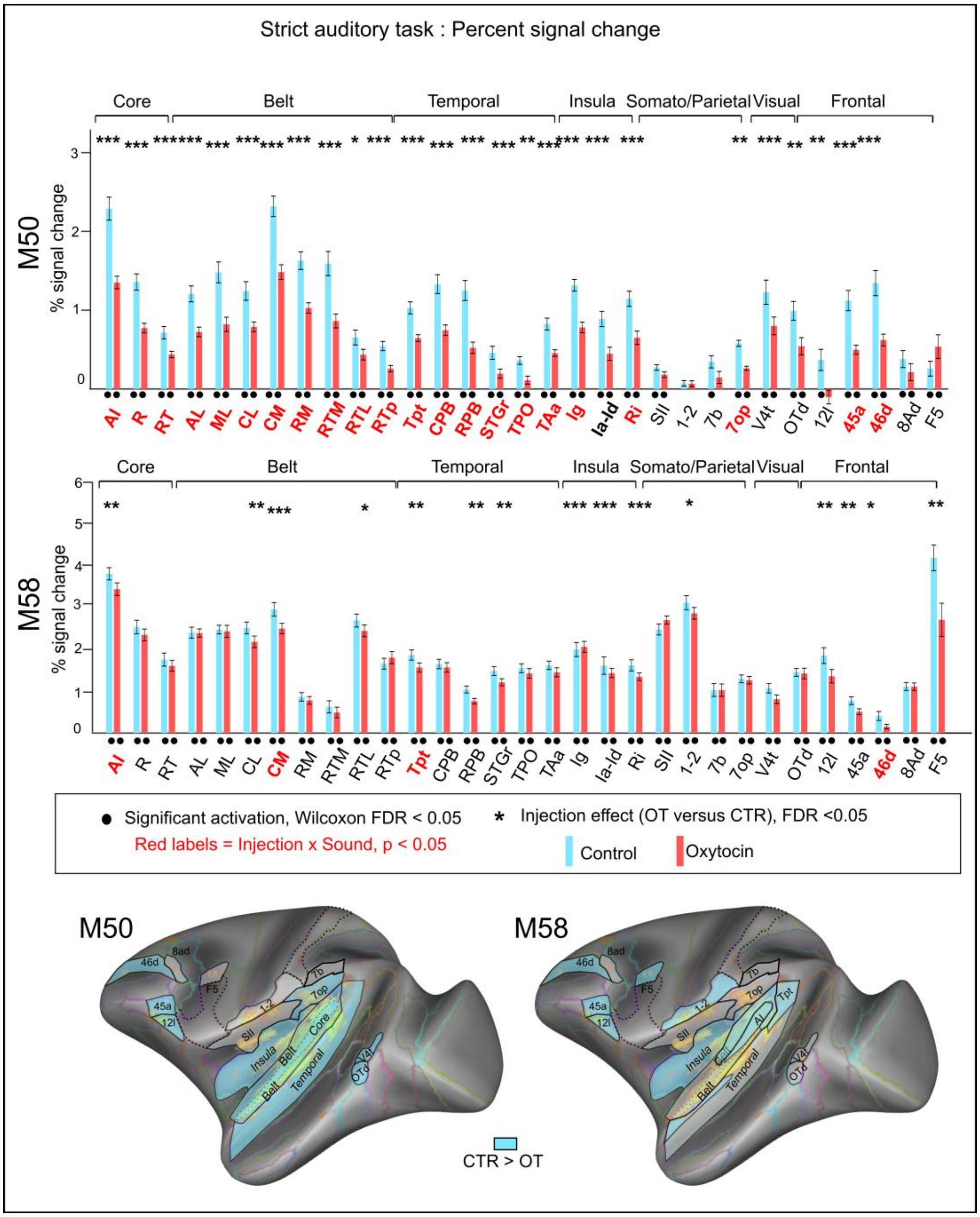


**Supplementary Figure 1. Bilateral auditory-responsive ROI analysis during the unimodal auditory task in each monkey.** Percent signal change (PSC) in functionally defined auditory-responsive cortical ROIs during the unimodal auditory task, shown separately for M50 and M58. Bars represent the mean PSC across the five intact sound categories (aggressive calls, coos, screams, animal sounds, and natural sounds) relative to fixation, for control (CTR, blue) and oxytocin (OT, red) sessions. For each ROI, values were averaged across homologous left and right ROIs. Error bars denote SEM across runs. ROIs are organized from early auditory regions (Core and Belt) to higher-order associative regions (temporal, insular, somato-parietal, visual, and frontal areas). Black dots below individual bars indicate responses significantly different from zero within each condition (Wilcoxon signed-rank tests, FDR-corrected at $p < 0.05$). Asterisks above paired bars indicate a significant main effect of Injection (ANOVA, FDR-corrected across ROIs, * $p<0.05$; ** $p<0.01$, *** $p<0.001$). Red bold ROI labels indicate regions showing an Injection × Sound interaction (uncorrected $p < 0.05$). The cortical insets provide schematic summaries of the bar plot results for each monkey, highlighting the ROIs showing stronger responses in CTR than OT sessions.

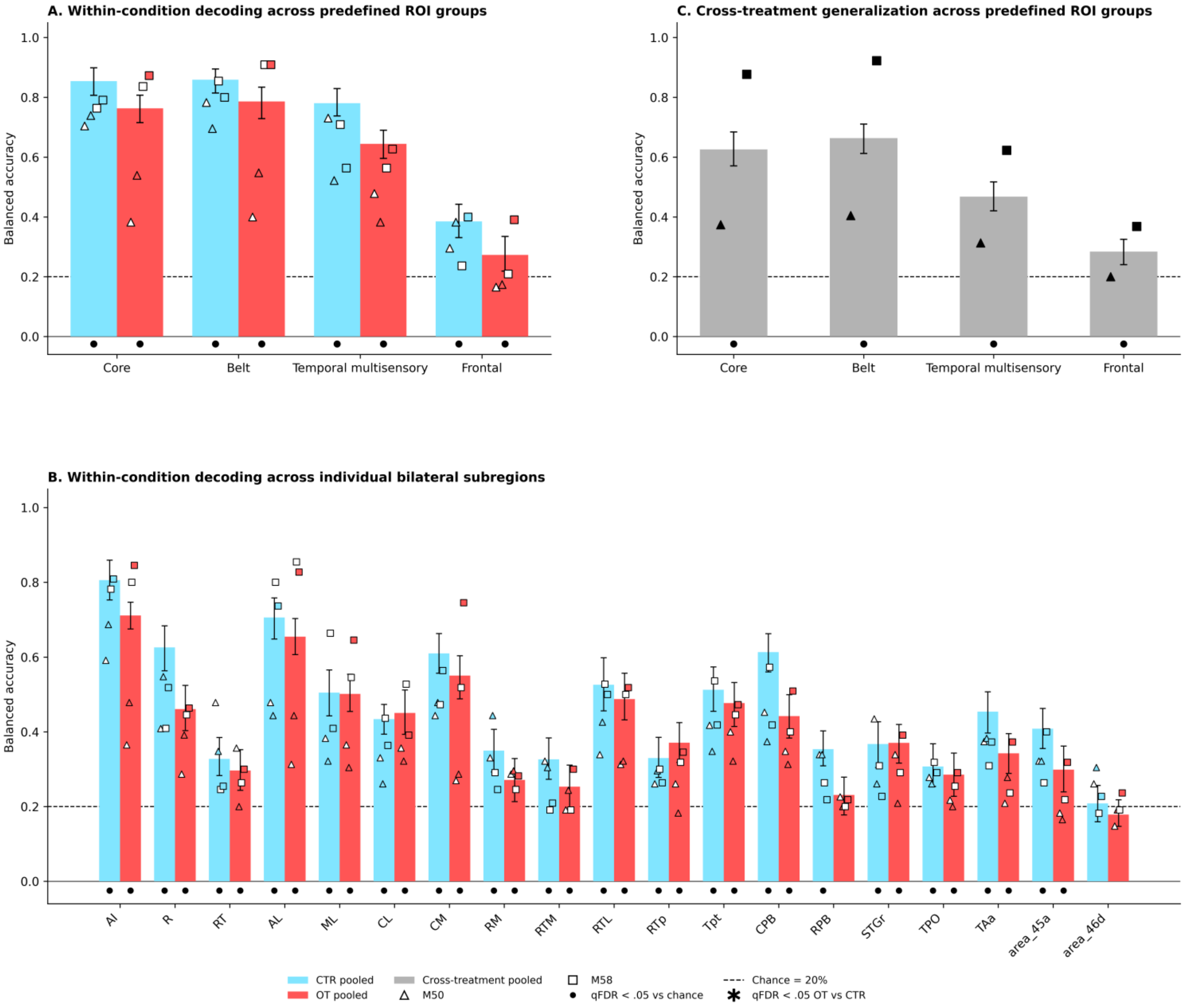


**Supplementary Figure 2. Within-condition and cross-treatment decoding of sound-category information in the unimodal auditory task. A)** Five-class multivariate decoding of aggressive calls, coos, screams, animal sounds and natural sounds within four predefined bilateral ROI groups (auditory Core, auditory Belt, temporal multisensory and frontal). Bars show pooled balanced accuracy under control (CTR; cyan) and oxytocin (OT; red), calculated as equal-weight means across M50 and M58. Error bars indicate 95% run-block bootstrap confidence intervals. Individual monkey-by-hemisphere estimates are overlaid (M50, triangles; M58, squares; open symbols, left hemisphere; filled symbols, right hemisphere). **B)**

Same analysis for the individual bilateral subregions comprising the four predefined ROI groups. Bars, error bars and individual symbols follow the conventions in A. **C)** Cross-treatment generalization of sound-category representations across the four predefined bilateral ROI groups. Classifiers were trained under CTR and tested under OT, and vice versa. Gray bars show the pooled bidirectional balanced accuracy, defined as the mean of CTR→OT and OT→CTR performance and averaged with equal weighting across monkeys. Error bars indicate 95% run-block bootstrap confidence intervals. Black triangles and squares show the corresponding bilateral bidirectional estimates for M50 and M58, respectively.

In all panels, the dashed horizontal line indicates the five-class chance level (20%). Black dots indicate pooled decoding significantly above chance after FDR correction ($q < 0.05$). Asterisks indicate significant OT–CTR differences in within-condition decoding after FDR correction of the session-level exact reassignment tests ($q < 0.05$). For cross-treatment decoding, significance against chance was assessed by permuting sound-category labels within training runs, retraining the classifier and testing on the unchanged opposite-treatment dataset, with FDR correction across the four predefined ROI groups.

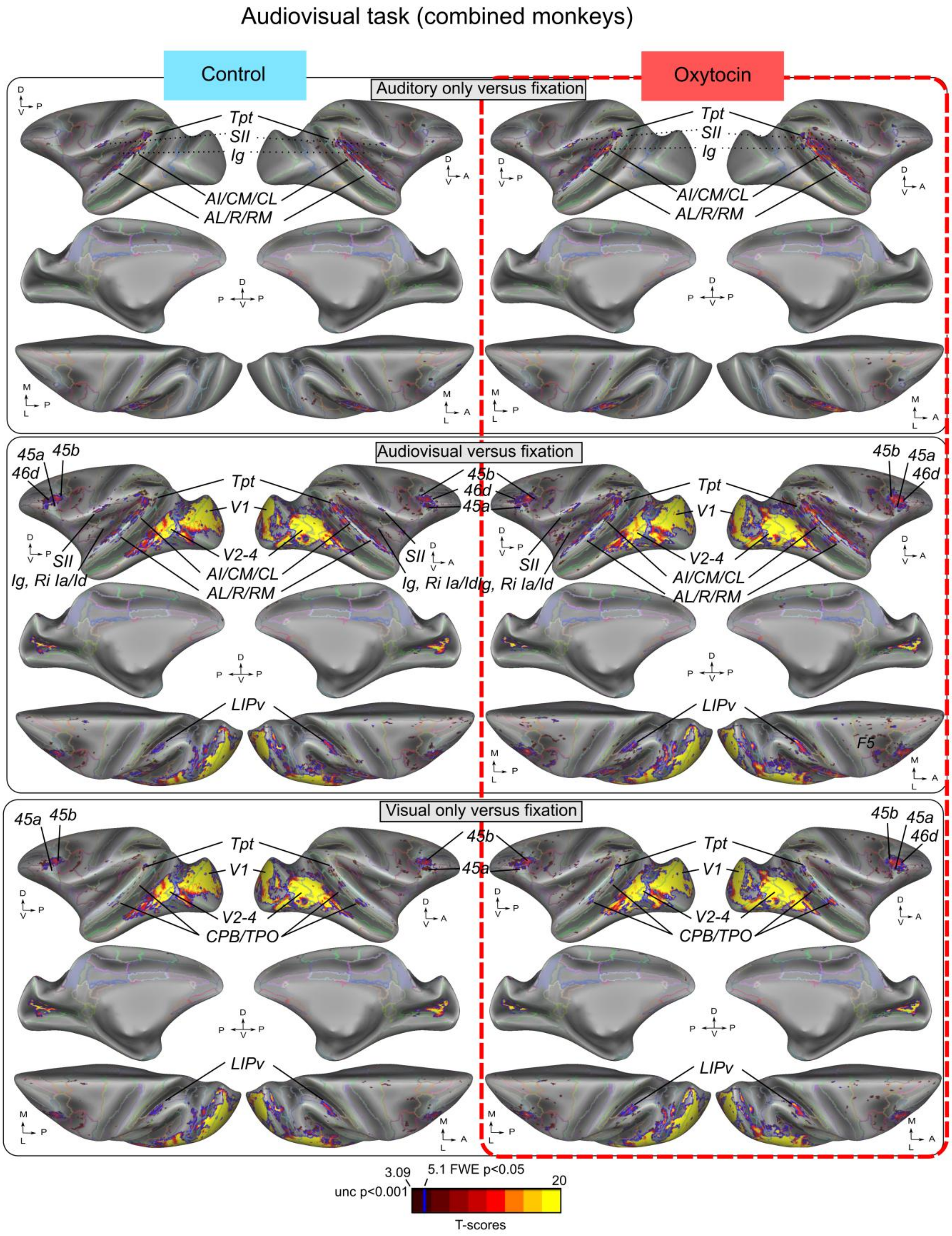


**Supplementary Figure 3. Task-evoked activation during the audiovisual task under control and oxytocin.** Surface projections show task-evoked activation maps for the auditory-only, audiovisual and visual-only conditions relative to fixation, displayed separately for control and oxytocin sessions in the combined-monkey analysis. Maps are projected onto inflated left and right MEBRAINS template surfaces. Warm colors indicate positive task-evoked responses relative to fixation. The color scale indicates *T*-scores, with reference thresholds shown below the maps.

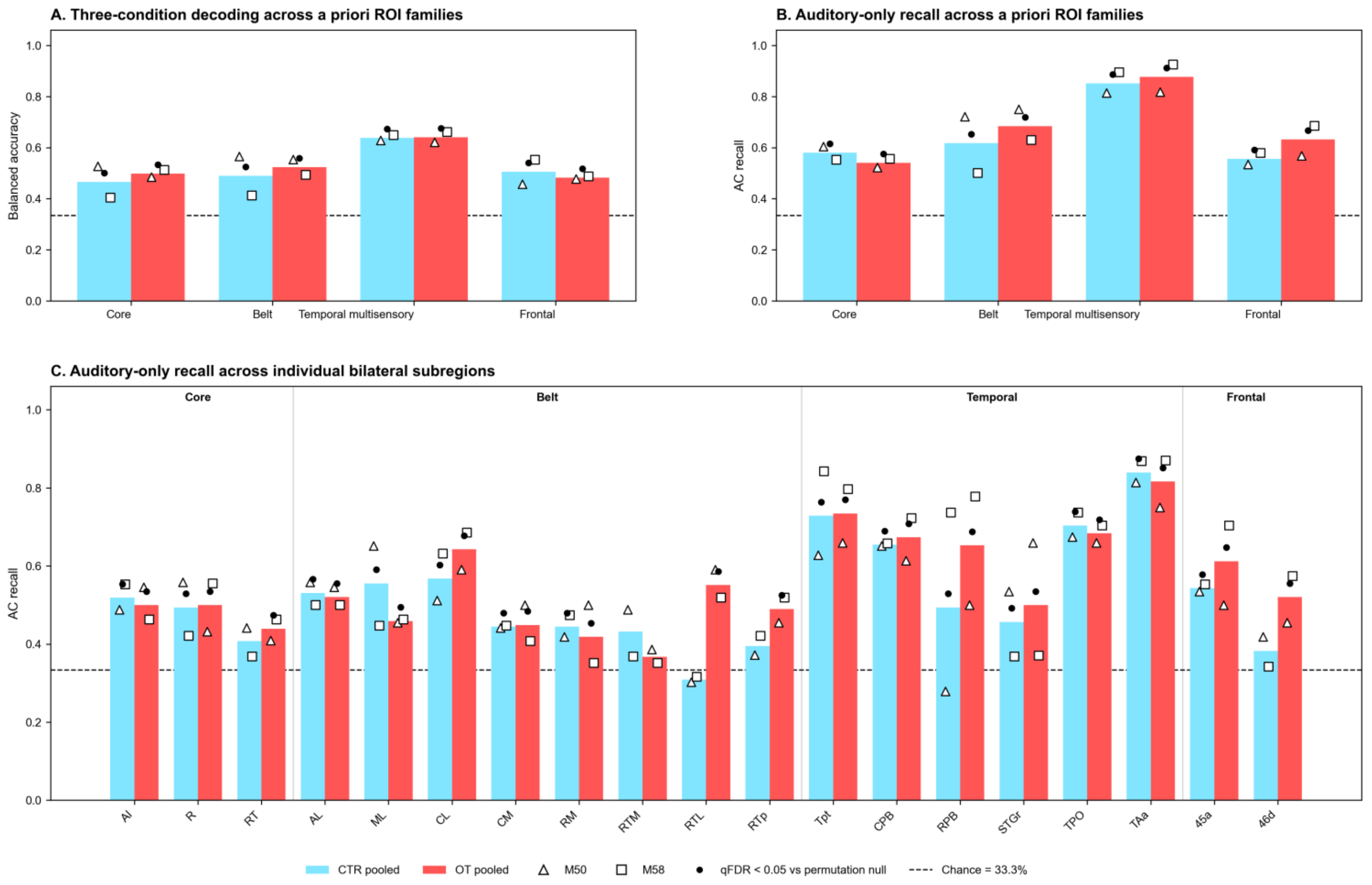


**Supplementary Figure 4. Multivariate decoding of auditory-only and audiovisual conditions during the audiovisual task. A)** Three-class decoding of auditory-only (AC), audiovisual-congruent (VAC) and audiovisual-incongruent (VAI) conditions within four bilateral a priori ROI families. **B)** Recall of the AC condition from the same three-class decoder within the four ROI families. **C)** AC recall across individual bilateral subregions, displayed in anatomical order from auditory Core and Belt regions to temporal and frontal association regions. Bars represent performance pooled equally across M50 and M58 for control (CTR, blue) and oxytocin (OT, red) sessions. Triangles and squares indicate bilateral decoding performance in M50 and M58, respectively. The dashed horizontal line indicates the 33.3% nominal chance level. Black dots indicate performance significantly above the permutation-derived null distribution after FDR correction across the relevant ROI set.

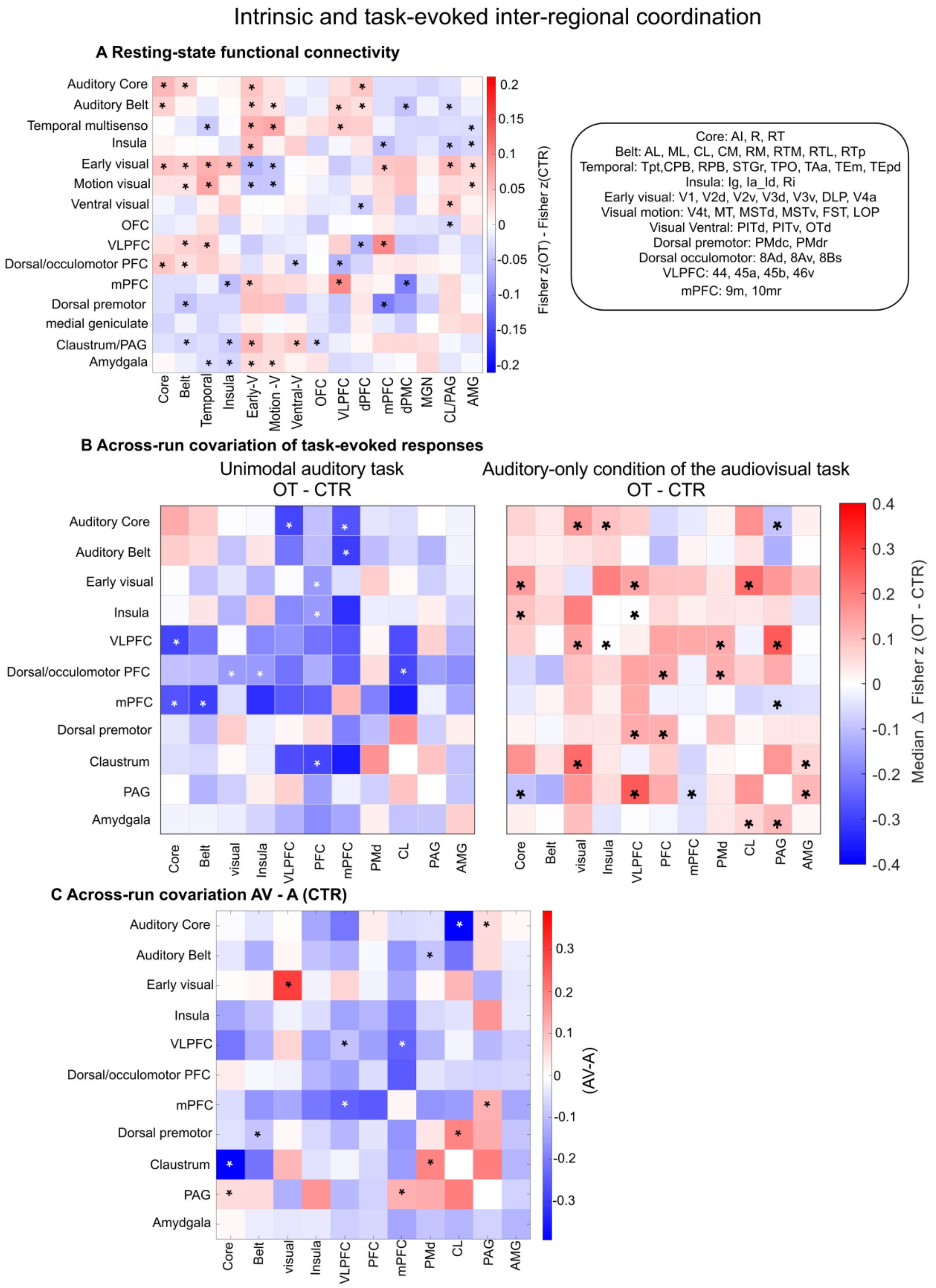


**Supplementary Figure 5. ROI-group matrices of intrinsic and task-evoked inter-regional coordination. A)** Resting-state functional connectivity differences between oxytocin (OT) and control (CTR) across predefined ROI groups. Each cell shows the OT-minus-CTR difference in mean Fisher-*z*-transformed connectivity for the corresponding ROI-group pair. Positive values indicate stronger connectivity under OT, while negative values indicated stronger connectivity under CTR. For each ROI-group pair, connectivity was first averaged across all constituent ROI-to-ROI edges within each run. OT–CTR effects were then estimated with equal weighting of the four monkey-by-hemisphere combinations (M50 left, M50 right, M58 left and M58 right). Asterisks indicate effects surviving BH-FDR correction ($q < 0.05$) across all unique testable ROI-group pairs in the complete matrix. Diagonal cells represent mean within-group connectivity across unique ROI pairs and not ROI self-correlations. **B)** Across-run covariation of task-evoked percent signal change (PSC) during the unimodal auditory task (left) and the auditory-only condition of the audiovisual task (right). Colors indicate the median OT-minus-CTR difference in Fisher-*z*-transformed across-run covariation across all ROI pairs belonging to each ROI-group pair. Positive values indicate relatively stronger/more positive covariation under OT and negative values relatively stronger/more positive covariation under CTR. Asterisks indicate ROI-group pairs containing at least one individual ROI-pair Injection effect surviving BH-FDR correction ($q < 0.05$) within the corresponding ROI-group pair. The median values shown by the color scale are descriptive and were not themselves subjected to a separate statistical test. **C)** Direct comparison of across-run PSC covariation between the auditory-only condition of the audiovisual task (AV) and the unimodal auditory task (A) under control conditions. Colors indicate the median AV-minus-A Fisher-*z* difference across ROI pairs within each ROI-group pair; positive values indicate relatively stronger/more positive covariation in AV and negative values relatively stronger/more positive covariation in A. Asterisks indicate ROI-group pairs containing at least one individual ROI-pair Task effect surviving BH-FDR correction ($q < 0.05$) within the corresponding ROI-group pair. The median values are descriptive and were not themselves subjected to a separate statistical test. Across-run covariation reflects covariation of task-evoked response amplitudes across runs and should not be interpreted as temporal functional connectivity. Matrices are symmetric because all relationships are undirected.

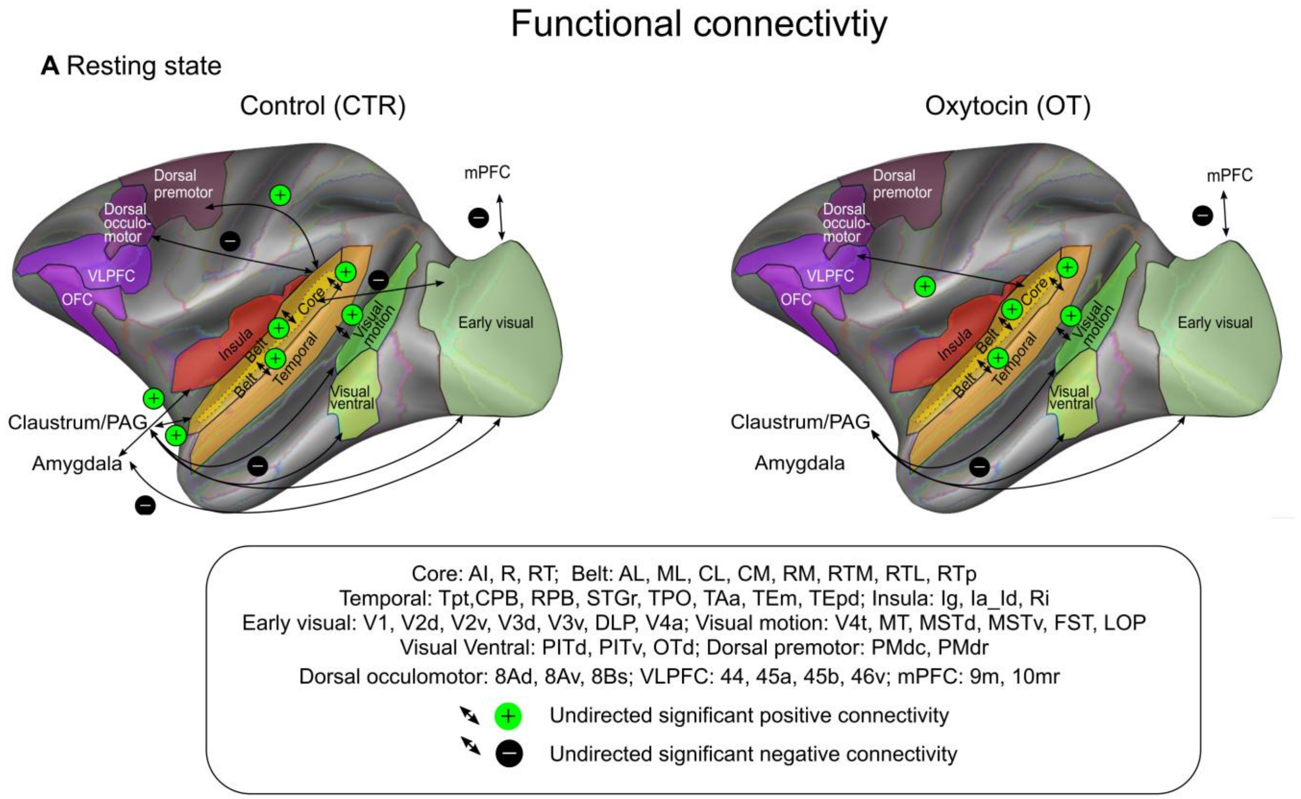


**Supplementary Figure 6: Resting-state functional connectivity under control and oxytocin.** Resting-state functional connectivity under control (CTR; left) and oxytocin (OT; right) across predefined connection families. Double-headed arrows denote undirected significant connectivity. Green plus and black minus symbols indicate significant positive and negative connectivity, respectively, after FDR correction.

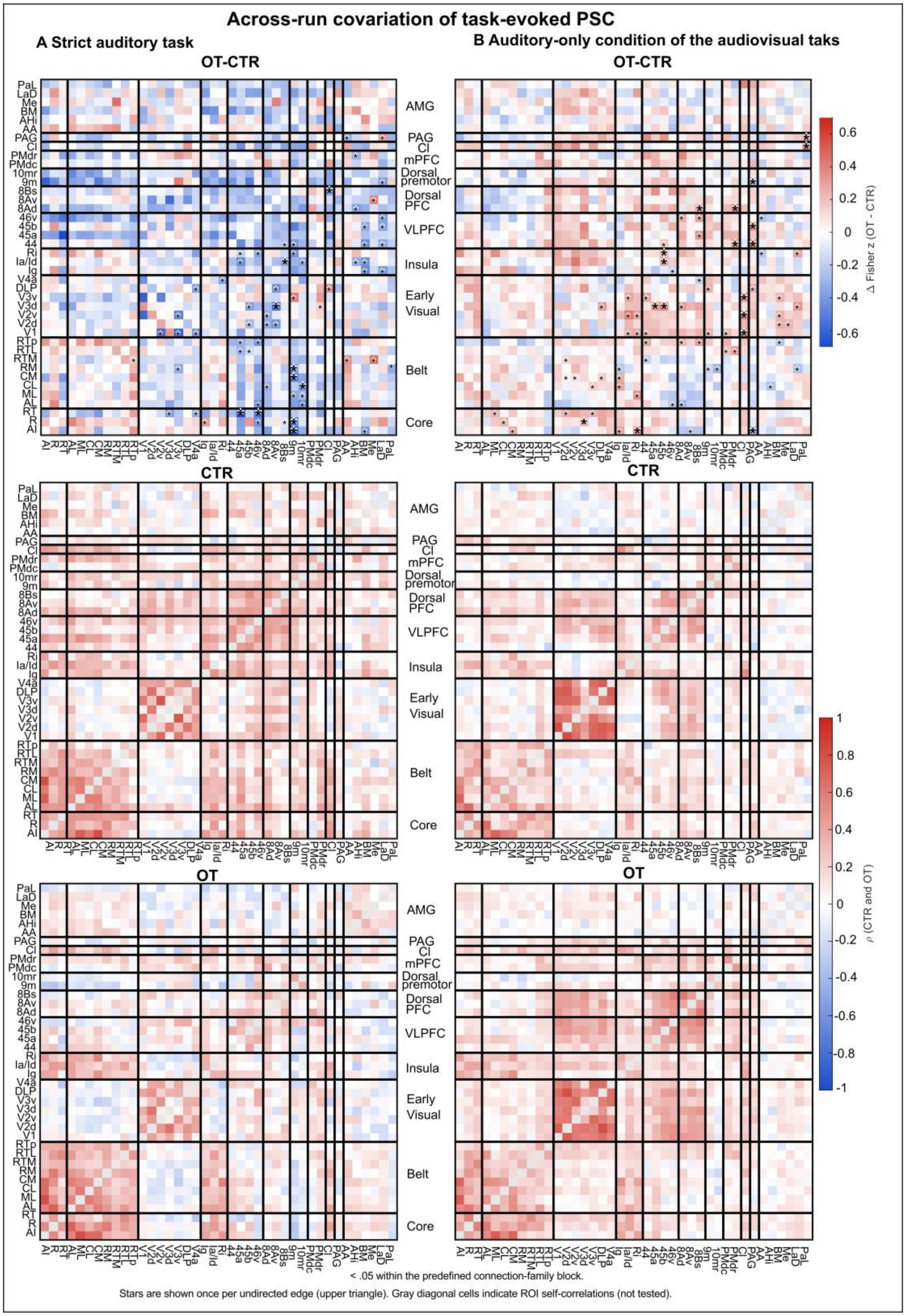


**Supplementary Figure 7: Across-run covariation of task-evoked PSC.** ROI-level PSC covariation during the unimodal auditory task (**A**) and the auditory-only condition of the audiovisual task (**B**). For each task, the top matrix shows the OT−CTR difference in Fisher-transformed Spearman correlations (Δ Fisher *z*), followed by the corresponding CTR and OT correlation matrices (ρ). ROIs are grouped by predefined connection families, delimited by black lines. In the OT−CTR matrices, asterisks indicate ROI-pair treatment effects at an uncorrected threshold of $p < 0.05$, with larger/bold asterisks indicating effects surviving FDR correction within the corresponding connection-family block.

### Supplementary tables

**Supplementary Table 1. Injection effects in auditory-responsive cortical ROIs across tasks.** A, C and E) Cross-monkey summaries of Injection effects. **B)** Complete pooled bilateral ROI-level statistics for the unimodal auditory task, including omnibus Injection, Injection × Sound, Injection × Hemisphere, and Injection × Monkey effects, together with sound-specific CTR–OT post-hoc comparisons. **D)** Complete pooled bilateral ROI-level statistics for the auditory-only condition of the audiovisual task. All cortical ROIs were functionally defined independently of the pharmacological comparison and anatomically constrained using RetCHARM.

***A. Unimodal auditory task***

| Analysis | Main Injection effect (FDR) | Direction among FDR ROIs | Left / Right ROIs | Overlap with pooled FDR ROIs | CTR > OT among pooled FDR ROIs | Injection × Sound interaction (FDR) |
|---|---|---|---|---|---|---|
| Bilateral ROIs | | | | | | |
| Pooled | 27/31 | 27 (all CTR > OT) | | | | 11/31 |
| M50 | 26/31 | 26 (all CTR > OT) | | 25/27 | 26/27 | 21/31 |
| M58 | 12/31 | 12 (all CTR > OT) | | 12/27 | 26/27 | 0/31 |
| Left/Right ROIs | | | | | | |
| Pooled | 42/71 | 1 OT > CTR; 41 CTR > OT | 29 / 13 | | | 13/71 |
| M50 | 49/71 | 49 (all CTR > OT) | 29 / 20 | 37/42 | 41/42 | 32/71 |
| M58 | 16/71 | 1 OT > CTR; 15 CTR > OT | 13 / 3 | 16/42 | 38/42 | 0/71 |

**Note.** Main Injection and Injection × Sound effects survived Benjamini–Hochberg FDR correction across the relevant ROI set. “Overlap with pooled FDR ROIs” indicates the number of pooled-significant ROIs also surviving FDR in the individual monkey. “CTR > OT among pooled FDR ROIs” indicates directional agreement irrespective of individual significance.

**B. Unimodal auditory task : complete pooled bilateral ROI statistics**

**Pooled: omnibus effects**

| ROI | Injection | Injection × Sound | Injection × Hemisphere | Injection × Monkey |
|---|---|---|---|---|
| AI | **F(1, 877)=91.14; p=1.29 × $10^{-20}$; q=3.99 × $10^{-19}$; $\eta p^2$=0.094** | **F(4, 877)=8.54; p=9.19 × $10^{-7}$; q=7.12 × $10^{-6}$; $\eta p^2$=0.037** | **F(1, 877)=20.54; p=6.64 × $10^{-6}$; q=6.87 × $10^{-5}$; $\eta p^2$=0.023** | **F(1, 877)=17.24; p=3.62 × $10^{-5}$; q=.00028; $\eta p^2$=0.019** |
| R | **F(1, 877)=32.93; p=1.31 × $10^{-8}$; q=4.53 × $10^{-8}$; $\eta p^2$=0.036** | F(4, 877)=2.50; p=.0412; q=.0852; $\eta p^2$=0.011 | **F(1, 877)=13.99; p=.00020; q=.0015; $\eta p^2$=0.016** | **F(1, 877)=8.56; p=.0035; q=.0091; $\eta p^2$=0.010** |
| RT | **F(1, 877)=7.54; p=.0062; q=.0076; $\eta p^2$=0.009** | F(4, 877)=2.17; p=.0702; q=.128; $\eta p^2$=0.010 | F(1, 877)=0.02; p=.8808; q=.9493; $\eta p^2$=0.000 | F(1, 877)=0.67; p=.412; q=.4912; $\eta p^2$=0.001 |
| AL | **F(1, 877)=22.68; p=2.24 × $10^{-6}$; q=5.34 × $10^{-6}$; $\eta p^2$=0.025** | **F(4, 877)=3.02; p=.0173; q=.0487; $\eta p^2$=0.014** | **F(1, 877)=12.55; p=.00042; q=.0026; $\eta p^2$=0.014** | **F(1, 877)=20.81; p=5.79 × $10^{-6}$; q=.00014; $\eta p^2$=0.023** |
| ML | **F(1, 877)=25.80; p=4.63 × $10^{-7}$; q=1.31 × $10^{-6}$; $\eta p^2$=0.029** | F(4, 877)=1.18; p=.3198; q=.4311; $\eta p^2$=0.005 | **F(1, 877)=20.94; p=5.42 × $10^{-6}$; q=6.87 × $10^{-5}$; $\eta p^2$=0.023** | **F(1, 877)=19.92; p=9.12 × $10^{-6}$; q=.00014; $\eta p^2$=0.022** |
| CL | **F(1, 877)=33.29; p=1.10 × $10^{-8}$; q=4.26 × $10^{-8}$; $\eta p^2$=0.037** | **F(4, 877)=4.51; p=.0013; q=.005; $\eta p^2$=0.020** | F(1, 877)=2.48; p=.1158; q=.1795; $\eta p^2$=0.003 | F(1, 877)=0.76; p=.3821; q=.4738; $\eta p^2$=0.001 |

| | | | | |
|---|---|---|---|---|
| CM | **F(1, 877)=85.03; p=2.13 × $10^{-19}$; q=3.30 × $10^{-18}$; $\eta p^2$=0.088** | **F(4, 877)=10.89; p=1.27 × $10^{-8}$; q=1.98 × $10^{-7}$; $\eta p^2$=0.047** | **F(1, 877)=9.47; p=.0022; q=.0083; $\eta p^2$=0.011** | **F(1, 877)=7.18; p=.0075; q=.0155; $\eta p^2$=0.008** |
| RM | **F(1, 877)=24.41; p=9.31 × $10^{-7}$; q=2.40 × $10^{-6}$; $\eta p^2$=0.027** | **F(4, 877)=4.11; p=.0026; q=.0081; $\eta p^2$=0.018** | F(1, 877)=4.06; p=.0442; q=.0806; $\eta p^2$=0.005 | **F(1, 877)=13.85; p=.00021; q=.00093; $\eta p^2$=0.016** |
| RTM | **F(1, 877)=19.73; p=1.01 × $10^{-5}$; q=1.95 × $10^{-5}$; $\eta p^2$=0.022** | F(4, 877)=0.83; p=.5045; q=.5852; $\eta p^2$=0.004 | F(1, 877)=0.06; p=.8039; q=.9493; $\eta p^2$=0.000 | **F(1, 877)=9.31; p=.0023; q=.0066; $\eta p^2$=0.011** |
| RTL | **F(1, 877)=11.98; p=.00057; q=.00097; $\eta p^2$=0.013** | **F(4, 877)=5.19; p=.00039; q=.0018; $\eta p^2$=0.023** | F(1, 877)=3.87; p=.0493; q=.085; $\eta p^2$=0.004 | F(1, 877)=0.05; p=.8248; q=.8816; $\eta p^2$=0.000 |
| RTp | F(1, 877)=0.83; p=.363; q=.363; $\eta p^2$=0.001 | F(4, 877)=1.13; p=.3394; q=.4384; $\eta p^2$=0.005 | F(1, 877)=0.01; p=.9079; q=.9493; $\eta p^2$=0.000 | **F(1, 877)=7.74; p=.0055; q=.0132; $\eta p^2$=0.009** |
| Tpt | **F(1, 877)=38.69; p=7.70 × $10^{-10}$; q=3.98 × $10^{-9}$; $\eta p^2$=0.042** | **F(4, 877)=8.88; p=5.00 × $10^{-7}$; q=5.17 × $10^{-6}$; $\eta p^2$=0.039** | **F(1, 877)=23.16; p=1.75 × $10^{-6}$; q=5.43 × $10^{-5}$; $\eta p^2$=0.026** | F(1, 877)=0.61; p=.4337; q=.4979; $\eta p^2$=0.001 |
| CPB | **F(1, 877)=33.59; p=9.46 × $10^{-9}$; q=4.19 × $10^{-8}$; $\eta p^2$=0.037** | F(4, 877)=2.88; p=.022; q=.0525; $\eta p^2$=0.013 | **F(1, 877)=7.68; p=.0057; q=.0161; $\eta p^2$=0.009** | **F(1, 877)=19.16; p=1.35 × $10^{-5}$; q=.00014; $\eta p^2$=0.021** |
| RPB | **F(1, 877)=80.31; p=1.90 × $10^{-18}$; q=1.96 × $10^{-17}$; $\eta p^2$=0.084** | F(4, 877)=1.45; p=.2154; q=.3179; $\eta p^2$=0.007 | F(1, 877)=0.01; p=.9186; q=.9493; $\eta p^2$=0.000 | **F(1, 877)=15.93; p=7.12 × $10^{-5}$; q=.00037; $\eta p^2$=0.018** |
| STGr | **F(1, 877)=21.77; p=3.55 × $10^{-6}$; q=7.86 × $10^{-6}$; $\eta p^2$=0.024** | F(4, 877)=2.02; p=.0903; q=.1473; $\eta p^2$=0.009 | **F(1, 877)=5.41; p=.0202; q=.0447; $\eta p^2$=0.006** | F(1, 877)=0.00; p=.9685; q=.9685; $\eta p^2$=0.000 |
| TPO | **F(1, 877)=9.75; p=.0019; q=.0028; $\eta p^2$=0.011** | F(4, 877)=2.09; p=.0803; q=.1382; $\eta p^2$=0.009 | **F(1, 877)=7.30; p=.007; q=.0181; $\eta p^2$=0.008** | F(1, 877)=1.10; p=.2937; q=.4108; $\eta p^2$=0.001 |
| TAa | **F(1, 877)=26.91; p=2.65 × $10^{-7}$; q=8.21 × $10^{-7}$; $\eta p^2$=0.030** | F(4, 877)=2.90; p=.0212; q=.0525; $\eta p^2$=0.013 | F(1, 877)=4.45; p=.0353; q=.0683; $\eta p^2$=0.005 | F(1, 877)=3.79; p=.0519; q=.0847; $\eta p^2$=0.004 |
| Ig | **F(1, 877)=8.07; p=.0046; q=.0062; $\eta p^2$=0.009** | **F(4, 877)=11.40; p=5.03 × $10^{-9}$; q=1.56 × $10^{-7}$; $\eta p^2$=0.049** | F(1, 877)=0.14; p=.7068; q=.8764; $\eta p^2$=0.000 | **F(1, 877)=13.14; p=.00031; q=.0012; $\eta p^2$=0.015** |
| Ia/Id | **F(1, 877)=9.13; p=.0026; q=.0037; $\eta p^2$=0.010** | F(4, 877)=2.78; p=.0258; q=.057; $\eta p^2$=0.013 | F(1, 877)=3.40; p=.0657; q=.1072; $\eta p^2$=0.004 | F(1, 877)=1.65; p=.199; q=.3002; $\eta p^2$=0.002 |
| Ri | **F(1, 877)=19.41; p=1.19 × $10^{-5}$; q=2.16 × $10^{-5}$; $\eta p^2$=0.022** | **F(4, 877)=5.17; p=.00041; q=.0018; $\eta p^2$=0.023** | F(1, 877)=0.24; p=.6276; q=.8107; $\eta p^2$=0.000 | F(1, 877)=1.62; p=.2034; q=.3002; $\eta p^2$=0.002 |
| SII | F(1, 877)=1.17; p=.2787; q=.3066; $\eta p^2$=0.001 | F(4, 877)=1.09; p=.3621; q=.449; $\eta p^2$=0.005 | **F(1, 877)=7.68; p=.0057; q=.0161; $\eta p^2$=0.009** | **F(1, 877)=6.59; p=.0104; q=.0202; $\eta p^2$=0.007** |
| areas 1-2 | **F(1, 877)=5.32; p=.0213; q=.0254; $\eta p^2$=0.006** | F(4, 877)=1.32; p=.2622; q=.3694; $\eta p^2$=0.006 | **F(1, 877)=9.65; p=.002; q=.0083; $\eta p^2$=0.011** | **F(1, 877)=4.96; p=.0262; q=.0478; $\eta p^2$=0.006** |
| area 7b | F(1, 877)=1.09; p=.2978; q=.3077; $\eta p^2$=0.001 | F(4, 877)=0.77; p=.5433; q=.6015; $\eta p^2$=0.004 | F(1, 877)=0.03; p=.8543; q=.9493; $\eta p^2$=0.000 | F(1, 877)=1.00; p=.3181; q=.4108; $\eta p^2$=0.001 |
| area 7op | **F(1, 877)=11.32; p=.00080; q=.0013; $\eta p^2$=0.013** | **F(4, 877)=6.36; p=4.78 × $10^{-5}$; q=.00030; $\eta p^2$=0.028** | F(1, 877)=0.00; p=.9906; q=.9906; $\eta p^2$=0.000 | **F(1, 877)=7.51; p=.0063; q=.0139; $\eta p^2$=0.008** |

| | | | | |
|---|---|---|---|---|
| V4t | **F(1, 877)=9.72; p=.0019; q=.0028; $\eta p^2$=0.011** | F(4, 877)=0.54; p=.7038; q=.7273; $\eta p^2$=0.002 | F(1, 877)=2.29; p=.1302; q=.1921; $\eta p^2$=0.003 | F(1, 877)=0.56; p=.4542; q=.5029; $\eta p^2$=0.001 |
| OTd | **F(1, 877)=4.98; p=.0259; q=.0298; $\eta p^2$=0.006** | F(4, 877)=0.49; p=.7449; q=.7449; $\eta p^2$=0.002 | **F(1, 877)=6.87; p=.0089; q=.0213; $\eta p^2$=0.008** | F(1, 877)=3.94; p=.0476; q=.082; $\eta p^2$=0.004 |
| area 12l | **F(1, 877)=21.03; p=$5.19 \times 10^{-6}$; q=$1.07 \times 10^{-5}$; $\eta p^2$=0.023** | F(4, 877)=2.25; p=.0617; q=.1196; $\eta p^2$=0.010 | F(1, 877)=1.01; p=.3146; q=.424; $\eta p^2$=0.001 | F(1, 877)=0.01; p=.909; q=.9393; $\eta p^2$=0.000 |
| area 45a | **F(1, 877)=58.18; p=$6.24 \times 10^{-14}$; q=$4.83 \times 10^{-13}$; $\eta p^2$=0.062** | F(4, 877)=1.88; p=.1122; q=.1739; $\eta p^2$=0.008 | F(1, 877)=5.09; p=.0243; q=.0502; $\eta p^2$=0.006 | **F(1, 877)=9.55; p=.0021; q=.0064; $\eta p^2$=0.011** |
| area 46d | **F(1, 877)=42.35; p=$1.28 \times 10^{-10}$; q=$7.95 \times 10^{-10}$; $\eta p^2$=0.046** | **F(4, 877)=4.25; p=.0021; q=.0071; $\eta p^2$=0.019** | **F(1, 877)=8.36; p=.0039; q=.0136; $\eta p^2$=0.009** | **F(1, 877)=9.58; p=.002; q=.0064; $\eta p^2$=0.011** |
| area 8ad | F(1, 877)=1.14; p=.2869; q=.3066; $\eta p^2$=0.001 | F(4, 877)=0.66; p=.6184; q=.661; $\eta p^2$=0.003 | F(1, 877)=1.75; p=.1862; q=.2624; $\eta p^2$=0.002 | F(1, 877)=1.01; p=.3156; q=.4108; $\eta p^2$=0.001 |
| F5 | **F(1, 877)=7.70; p=.0056; q=.0073; $\eta p^2$=0.009** | F(4, 877)=0.82; p=.5097; q=.5852; $\eta p^2$=0.004 | **F(1, 877)=10.79; p=.0011; q=.0055; $\eta p^2$=0.012** | **F(1, 877)=16.50; p=$5.31 \times 10^{-5}$; q=.00033; $\eta p^2$=0.018** |

**Pooled: sound-specific CTR–OT post-hoc comparisons**

| ROI | Aggressive calls | Coos | Screams | Animal sounds | Natural sounds |
|---|---|---|---|---|---|
| AI | **Δ=0.436; Z=2.46; p=.0141; q=.0176; r=0.366** | **Δ=1.293; Z=5.18; p=$2.27 \times 10^{-7}$; q=$1.14 \times 10^{-6}$; r=0.771** | **Δ=0.791; Z=4.26; p=$2.03 \times 10^{-5}$; q=$5.09 \times 10^{-5}$; r=0.635** | Δ=0.098; Z=1.15; p=.2519; q=.2519; r=0.171 | **Δ=0.649; Z=3.73; p=.00019; q=.00032; r=0.556** |
| R | Δ=0.186; Z=0.94; p=.3459; q=.3459; r=0.141 | **Δ=0.717; Z=3.34; p=.00085; q=.0043; r=0.497** | **Δ=0.456; Z=2.47; p=.0137; q=.0228; r=0.368** | Δ=0.132; Z=1.03; p=.3017; q=.3459; r=0.154 | **Δ=0.446; Z=3.06; p=.0022; q=.0054; r=0.457** |
| RT | Δ=0.052; Z=-0.03; p=.9775; q=.9775; r=-0.004 | **Δ=0.587; Z=3.15; p=.0016; q=.008; r=0.470** | Δ=-0.040; Z=-0.23; p=.817; q=.9775; r=-0.034 | Δ=0.118; Z=1.61; p=.1077; q=.1796; r=0.240 | Δ=0.346; Z=1.79; p=.0736; q=.1796; r=0.267 |
| AL | Δ=0.144; Z=1.20; p=.2293; q=.2866; r=0.179 | **Δ=0.587; Z=3.67; p=.00024; q=.0012; r=0.548** | Δ=0.074; Z=0.69; p=.4876; q=.4876; r=0.103 | Δ=0.205; Z=1.62; p=.1053; q=.2197; r=0.241 | Δ=0.235; Z=1.51; p=.1318; q=.2197; r=0.225 |
| ML | Δ=0.187; Z=0.82; p=.4132; q=.4132; r=0.122 | **Δ=0.548; Z=2.93; p=.0034; q=.0085; r=0.437** | Δ=0.245; Z=1.55; p=.1207; q=.1508; r=0.231 | Δ=0.276; Z=1.60; p=.1102; q=.1508; r=0.238 | **Δ=0.517; Z=3.04; p=.0024; q=.0085; r=0.453** |
| CL | Δ=0.125; Z=0.85; p=.3941; q=.4926; r=0.127 | **Δ=0.831; Z=4.55; p=$5.25 \times 10^{-6}$; q=$2.63 \times 10^{-5}$; r=0.679** | **Δ=0.611; Z=3.38; p=.00072; q=.0018; r=0.504** | Δ=0.079; Z=0.64; p=.5236; q=.5236; r=0.095 | Δ=0.318; Z=1.96; p=.0502; q=.0836; r=0.292 |
| CM | Δ=0.349; Z=1.81; p=.07; q=.0875; r=0.270 | **Δ=1.276; Z=4.95; p=$7.44 \times 10^{-7}$; q=$1.86 \times 10^{-6}$; r=0.738** | **Δ=1.085; Z=5.03; p=$4.94 \times 10^{-7}$; q=$1.86 \times 10^{-6}$; r=0.750** | Δ=0.067; Z=0.47; p=.6395; q=.6395; r=0.070 | **Δ=0.448; Z=3.39; p=.00069; q=.0012; r=0.506** |
| RM | Δ=0.260; Z=2.01; p=.0439; q=.0611; r=0.300 | **Δ=0.788; Z=3.79; p=.00015; q=.00076; r=0.565** | Δ=-0.078; Z=-0.27; p=.7908; q=.7908; r=-0.040 | **Δ=0.335; Z=2.48; p=.0132; q=.0331; r=0.369** | Δ=0.419; Z=1.97; p=.0489; q=.0611; r=0.294 |
| RTM | Δ=0.292; Z=0.83; p=.4067; q=.4132; r=0.124 | Δ=0.589; Z=2.57; p=.0102; q=.0512; r=0.383 | Δ=0.160; Z=0.82; p=.4132; q=.4132; r=0.122 | Δ=0.542; Z=2.00; p=.0451; q=.0752; r=0.299 | Δ=0.587; Z=2.20; p=.0281; q=.0703; r=0.327 |

| | | | | | |
|---|---|---|---|---|---|
| RTL | Δ=-0.017; Z=-0.42; p=.6721; q=.6721; r=-0.063 | **Δ=0.498; Z=2.97; p=.0029; q=.0147; r=0.443** | Δ=-0.241; Z=-1.31; p=.1885; q=.2356; r=-0.196 | **Δ=0.413; Z=2.51; p=.012; q=.02; r=0.374** | **Δ=0.497; Z=2.73; p=.0064; q=.016; r=0.406** |
| RTp | Δ=-0.032; Z=0.39; p=.697; q=.697; r=0.058 | Δ=0.261; Z=1.54; p=.1234; q=.2666; r=0.230 | Δ=-0.159; Z=-1.41; p=.1599; q=.2666; r=-0.209 | Δ=0.050; Z=0.57; p=.5687; q=.697; r=0.085 | Δ=0.256; Z=1.46; p=.1438; q=.2666; r=0.218 |
| Tpt | Δ=0.166; Z=1.47; p=.1407; q=.2217; r=0.220 | **Δ=0.840; Z=5.31; p=1.09 × $10^{-7}$; q=5.46 × $10^{-7}$; r=0.792** | **Δ=0.570; Z=3.81; p=.00014; q=.00035; r=0.568** | Δ=-0.064; Z=-0.86; p=.3879; q=.3879; r=-0.129 | Δ=0.180; Z=1.35; p=.1774; q=.2217; r=0.201 |
| CPB | Δ=0.035; Z=0.55; p=.5841; q=.5841; r=0.082 | **Δ=0.464; Z=3.53; p=.00042; q=.0021; r=0.526** | **Δ=0.468; Z=2.78; p=.0054; q=.009; r=0.415** | Δ=0.181; Z=1.57; p=.1153; q=.1442; r=0.235 | **Δ=0.531; Z=2.94; p=.0033; q=.0082; r=0.438** |
| RPB | Δ=0.297; Z=1.83; p=.0666; q=.0666; r=0.273 | **Δ=0.676; Z=4.86; p=1.18 × $10^{-6}$; q=5.89 × $10^{-6}$; r=0.724** | **Δ=0.424; Z=3.01; p=.0026; q=.0033; r=0.448** | **Δ=0.511; Z=3.54; p=.00040; q=.001; r=0.528** | **Δ=0.609; Z=3.01; p=.0026; q=.0033; r=0.448** |
| STGr | Δ=0.309; Z=1.55; p=.1207; q=.1508; r=0.231 | **Δ=0.368; Z=2.62; p=.0087; q=.0217; r=0.391** | Δ=-0.025; Z=-0.08; p=.9325; q=.9325; r=-0.013 | Δ=0.229; Z=1.76; p=.0792; q=.132; r=0.262 | **Δ=0.443; Z=3.30; p=.00096; q=.0048; r=0.492** |
| TPO | Δ=0.185; Z=1.87; p=.0617; q=.1029; r=0.278 | Δ=0.267; Z=2.17; p=.0298; q=.0745; r=0.324 | **Δ=0.386; Z=2.88; p=.0039; q=.0196; r=0.430** | Δ=-0.128; Z=-1.08; p=.2811; q=.2811; r=-0.161 | Δ=0.221; Z=1.71; p=.0873; q=.1091; r=0.255 |
| TAa | Δ=0.080; Z=0.59; p=.5535; q=.6918; r=0.088 | **Δ=0.509; Z=3.85; p=.00012; q=.00040; r=0.575** | **Δ=0.398; Z=3.78; p=.00016; q=.00040; r=0.563** | Δ=0.065; Z=0.24; p=.8083; q=.8083; r=0.036 | **Δ=0.288; Z=2.67; p=.0076; q=.0127; r=0.398** |
| Ig | Δ=-0.239; Z=-1.56; p=.118; q=.118; r=-0.233 | **Δ=0.916; Z=3.45; p=.00056; q=.0014; r=0.514** | **Δ=0.725; Z=3.46; p=.00054; q=.0014; r=0.516** | **Δ=-0.538; Z=-2.68; p=.0073; q=.0122; r=-0.400** | Δ=0.336; Z=1.71; p=.0873; q=.1091; r=0.255 |
| Ia/Id | Δ=0.136; Z=0.62; p=.5384; q=.5384; r=0.092 | **Δ=0.742; Z=2.83; p=.0047; q=.0235; r=0.422** | Δ=0.339; Z=1.55; p=.1207; q=.2011; r=0.231 | Δ=-0.230; Z=-1.13; p=.2566; q=.3208; r=-0.169 | Δ=0.560; Z=2.27; p=.0229; q=.0573; r=0.339 |
| Ri | Δ=0.111; Z=0.50; p=.6155; q=.6155; r=0.075 | **Δ=0.955; Z=3.89; p=.00010; q=.00050; r=0.580** | **Δ=0.527; Z=2.41; p=.016; q=.0266; r=0.359** | Δ=-0.193; Z=-0.93; p=.3517; q=.4397; r=-0.139 | **Δ=0.513; Z=2.57; p=.0102; q=.0256; r=0.383** |
| SII | Δ=-0.016; Z=-0.71; p=.4805; q=.6006; r=-0.105 | Δ=0.070; Z=0.99; p=.3233; q=.567; r=0.147 | Δ=-0.128; Z=-0.95; p=.3402; q=.567; r=-0.142 | Δ=-0.278; Z=-1.71; p=.0873; q=.4363; r=-0.255 | Δ=0.039; Z=0.38; p=.7053; q=.7053; r=0.056 |
| areas 1-2 | **Δ=0.320; Z=2.75; p=.006; q=.0299; r=0.410** | Δ=0.141; Z=1.03; p=.3017; q=.4848; r=0.154 | Δ=0.119; Z=1.04; p=.2964; q=.4848; r=0.156 | Δ=0.133; Z=0.86; p=.3879; q=.4848; r=0.129 | Δ=-0.083; Z=-0.36; p=.7222; q=.7222; r=-0.053 |
| area 7b | Δ=0.203; Z=0.21; p=.8346; q=.8346; r=0.031 | Δ=0.301; Z=1.43; p=.1533; q=.7662; r=0.213 | Δ=0.188; Z=0.74; p=.4597; q=.7662; r=0.110 | Δ=-0.011; Z=-0.27; p=.7908; q=.8346; r=-0.040 | Δ=-0.168; Z=-0.83; p=.4067; q=.7662; r=-0.124 |
| area 7op | Δ=0.026; Z=0.24; p=.8083; q=.8083; r=0.036 | **Δ=0.628; Z=4.33; p=1.50 × $10^{-5}$; q=7.50 × $10^{-5}$; r=0.645** | Δ=0.252; Z=1.86; p=.0633; q=.1583; r=0.277 | Δ=-0.140; Z=-1.51; p=.1318; q=.2197; r=-0.225 | Δ=0.114; Z=0.57; p=.5687; q=.7108; r=0.085 |
| V4t | Δ=0.051; Z=0.20; p=.8434; q=.8434; r=0.029 | Δ=0.427; Z=1.80; p=.0718; q=.1525; r=0.268 | Δ=0.475; Z=1.70; p=.0894; q=.1525; r=0.253 | Δ=0.475; Z=1.69; p=.0915; q=.1525; r=0.252 | Δ=0.288; Z=1.19; p=.2337; q=.2921; r=0.178 |
| OTd | Δ=0.263; Z=1.24; p=.2165; q=.3451; r=0.184 | Δ=0.225; Z=1.09; p=.276; q=.3451; r=0.162 | Δ=0.340; Z=1.35; p=.1774; q=.3451; r=0.201 | Δ=-0.028; Z=-0.28; p=.7821; q=.7821; r=-0.041 | Δ=0.400; Z=2.12; p=.0343; q=.1715; r=0.315 |
| area 12l | Δ=0.573; Z=2.09; p=.0363; q=.0907; r=0.312 | Δ=0.524; Z=1.90; p=.0572; q=.0953; r=0.284 | Δ=0.309; Z=1.64; p=.1005; q=.1256; r=0.245 | **Δ=0.967; Z=3.04; p=.0024; q=.0118; r=0.453** | Δ=0.018; Z=0.33; p=.7391; q=.7391; r=0.050 |

| | | | | | |
|---|---|---|---|---|---|
| area 45a | Δ=0.187; Z=1.63; p=.1029; q=.1029; r=0.243 | **Δ=0.613; Z=3.00; p=.0027; q=.0068; r=0.447** | **Δ=0.344; Z=2.40; p=.0165; q=.0206; r=0.358** | **Δ=0.497; Z=2.80; p=.005; q=.0084; r=0.418** | **Δ=0.575; Z=4.13; p=3.70 × $10^{-5}$; q=.00018; r=0.615** |
| area 46d | Δ=0.362; Z=1.69; p=.0915; q=.1525; r=0.252 | **Δ=0.737; Z=3.57; p=.00035; q=.001; r=0.533** | Δ=0.224; Z=1.29; p=.1962; q=.2337; r=0.193 | Δ=0.169; Z=1.19; p=.2337; q=.2337; r=0.178 | **Δ=0.961; Z=3.53; p=.00042; q=.001; r=0.526** |
| area 8ad | Δ=-0.113; Z=-0.75; p=.4529; q=.7548; r=-0.112 | Δ=0.275; Z=2.08; p=.0373; q=.1865; r=0.310 | Δ=0.037; Z=0.38; p=.7053; q=.8083; r=0.056 | Δ=0.157; Z=1.12; p=.2614; q=.6535; r=0.167 | Δ=0.074; Z=0.24; p=.8083; q=.8083; r=0.036 |
| F5 | Δ=1.071; Z=1.50; p=.1348; q=.3369; r=0.223 | Δ=0.975; Z=1.66; p=.0959; q=.3369; r=0.248 | Δ=0.046; Z=-0.52; p=.5997; q=.6234; r=-0.078 | Δ=0.311; Z=0.49; p=.6234; q=.6234; r=0.073 | Δ=0.486; Z=0.68; p=.4947; q=.6234; r=0.102 |

**Note.** Omnibus q values were FDR-corrected across the 31 bilateral ROIs separately for each effect. Sound-specific post-hoc comparisons used two-sided Wilcoxon signed-rank tests; q values were FDR-corrected across the five sounds within each ROI. All tests were included in the FDR correction; q values are displayed only for effects reaching the uncorrected threshold of p < .05. The pooled ANOVA included Monkey and the Injection × Monkey interaction. ηp² denotes partial eta squared.

***C. Auditory-only condition of the audiovisual task***

| Analysis | Main Injection effect, uncorrected | Direction, ROIs (uncorrected) | Main Injection effect, FDR | Direction, FDR ROIs | ROIs surviving FDR; (uncorrected only) |
|---|---|---|---|---|---|
| Bilateral ROIs | | | | | |
| Pooled | 10/31 | 9 OT > CTR; 1 CTR > OT | 9/31 | 8 OT > CTR; 1 CTR > OT | OT > CTR: AI, R, RT, AL, ML, CM, Ia/Id, F5; (RM). CTR > OT: area 46d |
| M50 | 10/31 | 3 OT > CTR; 7 CTR > OT | 5/31 | 2 OT > CTR; 3 CTR > OT | OT > CTR: AI, R; (ML). CTR > OT: Tpt, V4t, area 46d; (RTL, Ri, area 12l, area 8ad) |
| M58 | 16/31 | 16 (all OT > CTR) | 13/31 | 13 (all OT > CTR) | OT > CTR: R, RT, AL, CM, RTL, Tpt, TPO, TAa, Ri, area 7op, area 45a, area 8ad, F5; (CPB, RPB, Ia/Id) |
| Hemisphere-specific ROIs | | | | | |
| Pooled | 9/71 | 8 OT > CTR; 1 CTR > OT | 1/71 | 1 (all OT > CTR) | OT > CTR: right RT; (left AI, left R, left AL, left CM, right AI, right R, right ML). CTR > OT: (right area 46d) |
| M50 | 17/71 | 8 OT > CTR; 9 CTR > OT | 4/71 | 1 OT > CTR; 3 CTR > OT | OT > CTR: left AI; (left R, left RT, left AL, left ML, left CM, left RM, left F5). CTR > OT: left Tpt, left V4t, right area 46d; (left Ri, right CL, right RTL, right Tpt, right V4t, right area 45a) |
| M58 | 18/71 | 18 (all OT > CTR) | 1/71 | 1 (all OT > CTR) | OT > CTR: right RT; (left RTL, right AI, right R, right AL, right ML, right CL, right CM, right Tpt, right TPO, right TAa, right Ia/Id, right Ri, right area 7b, right area 7op, right area 45a, right area 8ad, right F5) |

**Note.** Uncorrected effects reached p < .05; FDR effects survived correction across the tested ROI set. ROIs outside parentheses survived FDR correction, whereas ROIs in parentheses were significant only at the uncorrected threshold.

**D. Auditory-only condition of the audiovisual task: pooled bilateral ROI statistics.** Values are run-level mean PSC ± SE. ΔPSC is defined as CTR − OT; negative values indicate larger responses under OT.

| ROI | CTR mean ± SE | OT mean ± SE | ΔPSC | Injection | Injection × Hemisphere | Injection × Monkey |
|---|---|---|---|---|---|---|

| | | | | | | |
|---|---|---|---|---|---|---|
| AI | 1.91 ± 0.11 | 2.38 ± 0.14 | -0.47 | **F(1, 317)=12.03; p=.00059; q=.0061; $\eta p^2$=0.037** | F(1, 317)=0.04; p=.8374; q=.9615; $\eta p^2$=0.000 | F(1, 317)=0.09; p=.7678; q=.8208; $\eta p^2$=0.000 |
| R | 1.22 ± 0.10 | 1.64 ± 0.11 | -0.41 | **F(1, 317)=13.27; p=.00031; q=.0049; $\eta p^2$=0.040** | F(1, 317)=0.00; p=.9498; q=.9814; $\eta p^2$=0.000 | F(1, 317)=0.39; p=.535; q=.6142; $\eta p^2$=0.001 |
| RT | 0.60 ± 0.09 | 1.05 ± 0.11 | -0.45 | **F(1, 317)=14.21; p=.00020; q=.0049; $\eta p^2$=0.043** | F(1, 317)=1.48; p=.2252; q=.8463; $\eta p^2$=0.005 | **F(1, 317)=6.73; p=.0099; q=.0305; $\eta p^2$=0.021** |
| AL | 1.36 ± 0.09 | 1.62 ± 0.10 | -0.26 | **F(1, 317)=8.21; p=.0045; q=.0312; $\eta p^2$=0.025** | F(1, 317)=0.66; p=.4165; q=.9384; $\eta p^2$=0.002 | F(1, 317)=3.66; p=.0565; q=.0876; $\eta p^2$=0.011 |
| ML | 1.16 ± 0.10 | 1.49 ± 0.10 | -0.33 | **F(1, 317)=7.64; p=.006; q=.0312; $\eta p^2$=0.024** | F(1, 317)=0.07; p=.7924; q=.9448; $\eta p^2$=0.000 | F(1, 317)=0.27; p=.6027; q=.6673; $\eta p^2$=0.001 |
| CL | 1.21 ± 0.08 | 1.25 ± 0.12 | -0.04 | F(1, 317)=0.26; p=.611; q=.7285; $\eta p^2$=0.001 | F(1, 317)=0.12; p=.7241; q=.9448; $\eta p^2$=0.000 | **F(1, 317)=5.54; p=.0191; q=.0432; $\eta p^2$=0.017** |
| CM | 1.88 ± 0.12 | 2.24 ± 0.13 | -0.37 | **F(1, 317)=7.93; p=.0052; q=.0312; $\eta p^2$=0.024** | F(1, 317)=1.39; p=.2393; q=.8463; $\eta p^2$=0.004 | **F(1, 317)=5.45; p=.0202; q=.0432; $\eta p^2$=0.017** |
| RM | 0.96 ± 0.12 | 1.21 ± 0.12 | -0.25 | F(1, 317)=4.36; p=.0376; q=.1166; $\eta p^2$=0.014 | F(1, 317)=0.10; p=.7565; q=.9448; $\eta p^2$=0.000 | F(1, 317)=0.00; p=.9884; q=.9884; $\eta p^2$=0.000 |
| RTM | 0.77 ± 0.13 | 0.89 ± 0.10 | -0.12 | F(1, 317)=0.78; p=.378; q=.558; $\eta p^2$=0.002 | F(1, 317)=0.54; p=.4646; q=.9448; $\eta p^2$=0.002 | F(1, 317)=3.21; p=.0742; q=.1096; $\eta p^2$=0.010 |
| RTL | 1.08 ± 0.11 | 1.12 ± 0.15 | -0.04 | F(1, 317)=0.31; p=.5794; q=.7184; $\eta p^2$=0.001 | F(1, 317)=2.15; p=.1438; q=.8463; $\eta p^2$=0.007 | **F(1, 317)=12.72; p=.00042; q=.0038; $\eta p^2$=0.039** |
| RTp | 0.68 ± 0.09 | 0.69 ± 0.11 | -0.01 | F(1, 317)=0.05; p=.8272; q=.8547; $\eta p^2$=0.000 | F(1, 317)=1.31; p=.2531; q=.8463; $\eta p^2$=0.004 | F(1, 317)=2.62; p=.1068; q=.1505; $\eta p^2$=0.008 |
| Tpt | 1.18 ± 0.08 | 1.09 ± 0.10 | 0.09 | F(1, 317)=0.42; p=.5192; q=.6706; $\eta p^2$=0.001 | F(1, 317)=1.15; p=.2842; q=.8463; $\eta p^2$=0.004 | **F(1, 317)=23.04; p=2.45 × $10^{-6}$; q=7.59 × $10^{-5}$; $\eta p^2$=0.068** |
| CPB | 0.97 ± 0.07 | 1.04 ± 0.08 | -0.07 | F(1, 317)=0.85; p=.3573; q=.5538; $\eta p^2$=0.003 | F(1, 317)=0.18; p=.6753; q=.9448; $\eta p^2$=0.001 | **F(1, 317)=6.57; p=.0108; q=.0305; $\eta p^2$=0.020** |
| RPB | 0.89 ± 0.09 | 0.91 ± 0.07 | -0.02 | F(1, 317)=0.12; p=.7262; q=.8026; $\eta p^2$=0.000 | F(1, 317)=0.08; p=.782; q=.9448; $\eta p^2$=0.000 | **F(1, 317)=5.39; p=.0209; q=.0432; $\eta p^2$=0.017** |
| STGr | 0.72 ± 0.08 | 0.57 ± 0.09 | 0.15 | F(1, 317)=2.33; p=.1279; q=.2832; $\eta p^2$=0.007 | F(1, 317)=0.22; p=.6379; q=.9448; $\eta p^2$=0.001 | F(1, 317)=0.93; p=.3344; q=.4147; $\eta p^2$=0.003 |
| TPO | 0.62 ± 0.07 | 0.70 ± 0.10 | -0.08 | F(1, 317)=1.19; p=.2771; q=.5054; $\eta p^2$=0.004 | F(1, 317)=0.02; p=.8959; q=.9652; $\eta p^2$=0.000 | **F(1, 317)=11.35; p=.00085; q=.0052; $\eta p^2$=0.035** |
| TAa | 0.81 ± 0.06 | 0.92 ± 0.08 | -0.11 | F(1, 317)=2.46; p=.1176; q=.2804; $\eta p^2$=0.008 | F(1, 317)=0.64; p=.4238; q=.9384; $\eta p^2$=0.002 | **F(1, 317)=10.74; p=.0012; q=.0058; $\eta p^2$=0.033** |
| Ig | 1.24 ± 0.14 | 1.38 ± 0.14 | -0.14 | F(1, 317)=1.04; p=.3087; q=.5317; $\eta p^2$=0.003 | F(1, 317)=0.09; p=.7696; q=.9448; $\eta p^2$=0.000 | F(1, 317)=4.71; p=.0307; q=.0595; $\eta p^2$=0.015 |

| | | | | | | |
|---|---|---|---|---|---|---|
| Ia/Id | 0.55 ± 0.10 | 0.92 ± 0.13 | -0.37 | **F(1, 317)=6.37; p=.0121; q=.0452; ηp²=0.020** | F(1, 317)=0.13; p=.7161; q=.9448; ηp²=0.000 | F(1, 317)=1.03; p=.3102; q=.4006; ηp²=0.003 |
| Ri | 1.29 ± 0.11 | 1.37 ± 0.13 | -0.08 | F(1, 317)=0.54; p=.4647; q=.6548; ηp²=0.002 | F(1, 317)=4.30; p=.0389; q=.603; ηp²=0.013 | **F(1, 317)=12.40; p=.00049; q=.0038; ηp²=0.038** |
| SII | 0.47 ± 0.08 | 0.59 ± 0.12 | -0.12 | F(1, 317)=1.51; p=.2201; q=.434; ηp²=0.005 | F(1, 317)=1.30; p=.2547; q=.8463; ηp²=0.004 | F(1, 317)=3.68; p=.0561; q=.0876; ηp²=0.011 |
| areas 1-2 | 0.65 ± 0.09 | 0.74 ± 0.12 | -0.10 | F(1, 317)=1.48; p=.224; q=.434; ηp²=0.005 | F(1, 317)=0.84; p=.3602; q=.9305; ηp²=0.003 | F(1, 317)=4.07; p=.0444; q=.081; ηp²=0.013 |
| area 7b | 0.25 ± 0.11 | 0.31 ± 0.10 | -0.06 | F(1, 317)=0.22; p=.6357; q=.7299; ηp²=0.001 | F(1, 317)=0.28; p=.5949; q=.9448; ηp²=0.001 | F(1, 317)=0.48; p=.4883; q=.5823; ηp²=0.002 |
| area 7op | 0.54 ± 0.07 | 0.66 ± 0.07 | -0.12 | F(1, 317)=2.86; p=.0919; q=.259; ηp²=0.009 | F(1, 317)=0.44; p=.5096; q=.9448; ηp²=0.001 | **F(1, 317)=7.63; p=.0061; q=.0209; ηp²=0.024** |
| V4t | 1.01 ± 0.18 | 0.66 ± 0.14 | 0.36 | F(1, 317)=2.56; p=.1109; q=.2804; ηp²=0.008 | F(1, 317)=0.01; p=.9029; q=.9652; ηp²=0.000 | **F(1, 317)=14.20; p=.00020; q=.003; ηp²=0.043** |
| OTd | 0.73 ± 0.11 | 0.68 ± 0.12 | 0.05 | F(1, 317)=0.10; p=.7508; q=.8026; ηp²=0.000 | F(1, 317)=1.35; p=.247; q=.8463; ηp²=0.004 | F(1, 317)=0.00; p=.968; q=.9884; ηp²=0.000 |
| area 12l | 0.80 ± 0.14 | 0.64 ± 0.19 | 0.16 | F(1, 317)=0.45; p=.503; q=.6706; ηp²=0.001 | F(1, 317)=0.00; p=.9917; q=.9917; ηp²=0.000 | **F(1, 317)=5.88; p=.0159; q=.041; ηp²=0.018** |
| area 45a | 0.57 ± 0.07 | 0.64 ± 0.08 | -0.06 | F(1, 317)=0.92; p=.337; q=.5498; ηp²=0.003 | F(1, 317)=1.08; p=.3003; q=.8463; ηp²=0.003 | **F(1, 317)=10.52; p=.0013; q=.0058; ηp²=0.032** |
| area 46d | 0.75 ± 0.122 | 0.41 ± 0.11 | 0.344 | **F(1, 317)=6.22; p=.0131; q=.0452; ηp²=0.019** | F(1, 317)=4.45; p=.0358; q=.603; ηp²=0.014 | F(1, 317)=2.49; p=.1155; q=.1556; ηp²=0.008 |
| area 8ad | 0.75 ± 0.10 | 0.71 ± 0.11 | 0.045 | F(1, 317)=0.02; p=.8811; q=.8811; ηp²=0.000 | F(1, 317)=2.02; p=.1559; q=.8463; ηp²=0.006 | **F(1, 317)=9.43; p=.0023; q=.009; ηp²=0.029** |
| F5 | 0.68 ± 0.24 | 1.54 ± 0.29 | -0.864 | **F(1, 317)=6.48; p=.0114; q=.0452; ηp²=0.020** | F(1, 317)=0.07; p=.7886; q=.9448; ηp²=0.000 | F(1, 317)=3.76; p=.0535; q=.0876; ηp²=0.012 |

**Note.** Omnibus q values were FDR-corrected across the 31 bilateral ROIs separately for each effect. All tests were included in the FDR correction; q values are displayed only for effects reaching the uncorrected threshold of $p < .05$. The pooled ANOVA included Monkey and the Injection × Monkey interaction. ηp² denotes partial eta squared.

**E. Visual-only condition of the audiovisual task: cross-monkey summary of Injection effects**

| Analysis | Main Injection effect, uncorrected | Direction, ROIs (uncorrected) | Main Injection effect, FDR | Direction, FDR ROIs | ROIs surviving FDR; (uncorrected only) |
|---|---|---|---|---|---|
| Bilateral | | | | | |
| Pooled | 3/31 | 2 OT > CTR; 1 CTR > OT | 0/31 | 0 | OT > CTR: (area 7op, V4t). CTR > OT: (STGr) |
| M50 | 8/31 | 8 (all CTR > OT) | 0/31 | 0 | CTR > OT: (CL, RTL, RTp, Tpt, STGr, area 12l, area 45a, F5) |

| | | | | | |
|---|---|---|---|---|---|
| M58 | 4/31 | 4 (all OT > CTR) | 3/31 | 3 (all OT > CTR) | OT > CTR: area 7op, V4t, area 45a; (CM) |
| Hemisphere-specific ROIs | | | | | |
| Pooled | 5/71 | 2 OT > CTR; 3 CTR > OT | 0/71 | 0 | OT > CTR: (left TAa, right area 7op). CTR > OT: (left area 12l, right RTL, right areas 1-2) |
| M50 | 9/71 | 1 OT > CTR; 8 CTR > OT | 0/71 | 0 | OT > CTR: (right area 7op). CTR > OT: (left Tpt, left MT, left area 12l, right CL, right RTL, right RPB, right STGr, right area 45a) |
| M58 | 9/71 | 9 (all OT > CTR) | 0/71 | 0 | OT > CTR: (left TPO, left TAa, left area 7a, left area 7op, left V4t, left area 45a, right RT, right CM, right area 45a) |

*Note. Uncorrected effects reached p < .05; FDR effects survived correction across the tested ROI set. ROIs outside parentheses survived FDR correction, whereas ROIs in parentheses were significant only at the uncorrected threshold.*

**Supplementary Table 2A. Auditory responsiveness and Injection effects in pooled auditory-responsive subcortical ROIs during the unimodal auditory task.** Rows are restricted to the 17 SARM ROIs that met the pooled exploratory auditory-response criterion (CTR and OT collapsed across all 10 sound conditions; two-sided Wilcoxon signed-rank test, nominal p < .05, and positive median PSC). For each analysis, the Auditory response column reports the median condition-collapsed PSC, Wilcoxon Z statistic, raw p value, and Benjamini-Hochberg FDR-adjusted q value; screening q values were corrected across all 50 SARM ROIs. Injection effects were tested only for ROIs that met the corresponding analysis-specific auditory-response criterion. To keep the table compact, Injection effect columns report only nominal effects (p < .05), including direction, F statistic, raw p value, FDR-adjusted q value, and partial eta-squared. A dash (—) indicates that the Injection test was performed but did not reach nominal significance; NR indicates that the ROI did not meet the auditory-response criterion, and NT indicates that the Injection effect was therefore not tested. Injection q values were corrected across the analysis-specific responsive set (Pooled: 17 ROIs; M50: 10 ROIs; M58: 18 ROIs). Bold q values indicate Injection effects surviving FDR correction at q < .05. ROIs meeting an individual-monkey auditory-response criterion but not the pooled criterion are reported in Supplementary Table 2B.

| ROI | Pooled | | M50 | | M58 | |
|---|---|---|---|---|---|---|
| | Auditory response (Median PSC; Z; p; q) | Injection effect (Direction; F; p; q; ηp²) | Auditory response (Median PSC; Z; p; q) | Injection effect (Direction; F; p; q; ηp²) | Auditory response (Median PSC; Z; p; q) | Injection effect (Direction; F; p; q; ηp²) |
| **AH_R** | Mdn = 0.189; Z = 2.51 p = .012; q = .043 | OT > CTR F(1, 880) = 4.13; p = .042; q = .180; ηp² = 0.005 | *NR* | *NT* | Mdn = 0.406; Z = 2.38 p = .017; q = .059 | — |
| **AHi_R** | Mdn = 0.060; Z = 2.34; p = .019; q = .064 | — | *NR* | *NT* | Mdn = 0.470; Z = 2.79 p = .005; q = .024 | — |
| **APul_L** | Mdn = 0.139; Z = 3.26; p = .001; q = .007 | — | Mdn = 0.139; Z = 2.90 p = .004; q = .044 | — | *NR* | *NT* |
| **APul_R** | Mdn = 0.157; Z = 3.56; p < .001; q = .003 | — | Mdn = 0.182; Z = 3.58 p < .001; q = .009 | — | *NR* | *NT* |
| **BLD_L** | Mdn = 0.136; Z = 3.00; p = .003; q = .015 | — | *NR* | *NT* | Mdn = 0.249; Z = 2.84 | — |

| | | | | | | |
|---|---|---|---|---|---|---|
| | | | | | p = .005; q = .023 | |
| **BLV_L** | Mdn = 0.075; Z = 2.20 p = .028; q = .083 | — | Mdn = 0.116; Z = 1.98 p = .047; q = .197 | — | *NR* | *NT* |
| **BM_L** | Mdn = 0.124; Z = 2.72 p = .007; q = .027 | — | *NR* | *NT* | Mdn = 0.115; Z = 2.15 p = .032; q = .088 | — |
| **CI_L** | Mdn = 0.054; Z = 4.09 p < .001; q < .001 | CTR > OT F(1, 880) = 8.37; p = .004; q **= .044**; ηp² = 0.009 | Mdn = 0.033; Z = 2.41 p = .016; q = .080 | — | Mdn = 0.116; Z = 3.24 p = .001; q = .010 | CTR > OT F(1, 420) = 6.61; p = .010; q = .109; ηp² = 0.016 |
| **CI_R** | Mdn = 0.047; Z = 3.66 p < .001; q = .002 | CTR > OT F(1, 880) = 7.87; p = .005; q **= .044**; ηp² = 0.009 | Mdn = 0.052; Z = 4.18 p < .001; q = .001 | — | *NR* | *NT* |
| **LPul_L** | Mdn = 0.097; Z = 2.75 p = .006; q = .027 | — | *NR* | *NT* | Mdn = 0.212; Z = 2.89 p = .004; q = .023 | — |
| **MGN_L** | Mdn = 0.548; Z = 5.99 p < .001; q < .001 | — | Mdn = 0.275; Z = 3.13, p = .002; q = .029 | — | Mdn = 1.115; Z = 4.82 p < .001; q < .001 | OT > CTR F(1, 420) = 6.35; p = .012; q = .109; ηp² = 0.015 |
| **MGN_R** | Mdn = 0.493; Z = 5.49 p < .001; q < .001 | — | Mdn = 0.286; Z = 2.85 p = .004; q = .044 | CTR > OT F(1, 440) = 11.63; p < .001; q **= .007**; ηp² = 0.026 | Mdn = 0.734; Z = 4.67 p < .001; q < .001 | — |
| **MPul_L** | Mdn = 0.184; Z = 2.86 p = .004; q = .021 | — | Mdn = 0.192; Z = 2.63, p = .009; q = .061 | — | *NR* | *NT* |
| **PAG_L** | Mdn = 0.278; Z = 4.74 p < .001; q < .001 | — | Mdn = 0.147; Z = 2.41, p = .016; q = .080 | CTR > OT F(1, 440) = 8.07; p = .005; q **= .024**; ηp² = 0.018 | Mdn = 0.329; Z = 4.17, p < .001; q < .001 | — |
| **PAG_R** | Mdn = 0.294; Z = 4.99 p < .001; q < .001 | CTR > OT F(1, 880) = 4.60; p = .032; | Mdn = 0.218; Z = 2.47 p = .013; q = .080 | CTR > OT F(1, 440) = 6.07; p = .014; q | Mdn = 0.375; Z = 4.55 p < .001; q < .001 | — |

| | | | | | | |
|---|---|---|---|---|---|---|
| | | q = .180; $\eta p^2$ = 0.005 | | **= .047**; $\eta p^2$ = 0.014 | | |
| **PaL_L** | Mdn = 0.135;<br>Z = 2.19 p = .028;<br>q = .083 | — | *NR* | *NT* | *NR* | *NT* |
| **STIA_L** | Mdn = 0.178;<br>Z = 2.60 p = .009;<br>q = .036 | — | *NR* | *NT* | Mdn = 0.516; Z = 2.91<br>p = .004; q = .023 | — |

Abbreviations: AH, anterior hypothalamic area; AHi, amygdalohippocampal area; APul, anterior pulvinar; BLD, basolateral dorsal amygdaloid nucleus; BLV, basolateral ventral amygdaloid nucleus; BM, basomedial amygdaloid nucleus; Cl, claustrum; LPul, lateral pulvinar; MGN, medial geniculate nucleus; MPul, medial pulvinar; PAG, periaqueductal grey; PaL, paralaminar amygdaloid nucleus; STIA, intra-amygdaloid stria terminalis; NR, not responsive; NT, not tested.

**Supplementary Table 2B. Additional M58-specific auditory-responsive subcortical ROIs during the unimodal auditory task.** Seven additional SARM ROIs met the exploratory auditory-response criterion in M58 (two-sided Wilcoxon signed-rank $p < .05$ with positive median PSC) but not in the pooled analysis. No M50-specific ROIs were identified: all 10 M50-responsive ROIs were already included in the pooled 17-ROI set shown in Supplementary Table 2A. Auditory-screen q values were Benjamini-Hochberg FDR-corrected across all 50 SARM ROIs. Injection statistics are reported for completeness and were FDR-corrected across the 18 M58 auditory-responsive ROIs. ΔPSC is defined as CTR − OT; positive values indicate CTR > OT. None of these additional ROIs showed a nominal Injection effect (all $p \geq .095$).

Abbreviations: AA, anterior amygdaloid area; ASt, amygdalostriatal transition area; BLI, basolateral intermediate amygdaloid nucleus; Ce, central amygdaloid nucleus; DEn, dorsal endopiriform claustrum; EA, extended amygdala; LPul, lateral pulvinar.

| ROI | M58 auditory screen | M58 Injection effect |
|---|---|---|
| AA_L | Median PSC=0.261; Z=2.49; p=.013; q=.0497 | ΔPSC=0.168 (CTR > OT); $F(1,420)$=2.16; p=.142; q=.365; $\eta p^2$=0.005 |
| ASt_L | Median PSC=0.284; Z=2.71; p=.007; q=.028 | ΔPSC=0.189 (CTR > OT); $F(1,420)$=1.72; p=.191; q=.419; $\eta p^2$=0.004 |
| BLI_R | Median PSC=0.263; Z=2.37; p=.018; q=.059 | ΔPSC=-0.034 (OT > CTR); $F(1,420)$=0.05; p=.822; q=.934; $\eta p^2$=0.000 |
| Ce_R | Median PSC=0.410; Z=2.26; p=.024; q=.069 | ΔPSC=-0.224 (OT > CTR); $F(1,420)$=1.58; p=.209; q=.419; $\eta p^2$=0.004 |
| DEn_R | Median PSC=0.298; Z=2.85; p=.004; q=.023 | ΔPSC=0.010 (CTR > OT); $F(1,420)$=0.00; p=.945; q=.945; $\eta p^2$=0.000 |
| EA_L | Median PSC=0.240; Z=2.30; p=.022; q=.067 | ΔPSC=0.059 (CTR > OT); $F(1,420)$=0.08; p=.779; q=.934; $\eta p^2$=0.000 |
| LPul_R | Median PSC=0.349; Z=3.91; p=$9.2 \times 10^{-5}$; q=$9.2 \times 10^{-4}$ | ΔPSC=-0.217 (OT > CTR); $F(1,420)$=2.81; p=.095; q=.288; $\eta p^2$=0.007 |

**Supplementary Table 3**. **Cortical and subcortical ROI sets used for percent-signal-change analyses.** Cortical ROIs were defined independently of the pharmacological comparison using pooled CTR and OT data and constrained by RetCHARM parcels. Auditory cortical ROIs were selected from the unimodal auditory task, visual cortical ROIs from visual stimulation in the audiovisual task, and subcortical ROIs were anatomically defined using SARM. Homologous cortical ROIs were combined for bilateral summaries, with hemisphere-specific analyses performed where indicated. The subcortical set comprised 25 bilateral structures (50 hemispheric ROIs).

| ROI set | ROI labels included | Definition |
|---|---|---|
| Auditory cortical ROIs | Bilateral homologous set (31):<br>Core: AI, R, RT;<br>Belt: AL, ML, CL, CM, RM, RTM, RTL, RTp;<br>Temporal/multisensory: Tpt, CPB, RPB, STGr, TPO, TAa;<br>Insula: Ig, Ia/Id, Ri;<br>Somato/parietal: SII, areas 1-2, 7b, 7op;<br>Visual-related: V4t, OTd;<br>Frontal: area 12l, area 45a, area 46d, area 8ad, F5.<br>Additional hemisphere-specific ROIs: left TEm, TEpd, area 7a, MT, V4a, PITd; right TGdg, PGa, MSTd. | Pooled CTR + OT unimodal auditory data (All sounds > Silence, $P_{\_FWE} < 0.001$), constrained by RetCHARM parcels |
| Visual cortical ROIs | Striate/extrastriate: V1, V2d, V3d, DLP, V3v, V2v, V4, V4a<br>Motion/dorsal: V4t, MT, MSTd<br>Ventral temporal: PITd, PITv, OTd, TEm<br>Parietal/visuomotor: LIPv, LIPd, area 7a, area 7op<br>Temporal/multisensory: CPB, RPB, Tpt, TPO, TAa<br>Frontal: areas 8Ad, 8Av, 8Bs, 44, 45a, 45b, 46v, 12m, 12r, 13l, F5, PMdc, preSMA<br>Somato/motor: area 3a-b | Pooled CTR + OT audiovisual-task data (Visual stimulation > Fixation, $P_{\_FWE} <$ 0.001), constrained by RetCHARM parcels |
| Subcortical SARM ROIs | Bilateral structures: AA, AHi, AH, APir, APul, ASt, BLD, BLI, BLV, BM, Ce, Cl, DEn, EA, IPul, LaD, LaV, LPul, Me, MG, MPul, PAG, PaL, STIA, VCo | Anatomically defined using SARM |

**Supplementary Table 4. ROI masks and aggregation levels used in the unimodal auditory multivoxel pattern analysis.** The table lists the ROI masks used for five-class decoding of aggressive calls, coos, screams, animal sounds and natural sounds. Analyses were performed separately in left and right hemispheres and in bilateral unions where indicated.

| ROI level | Masks included | Hemisphere aggregation |
|---|---|---|
| Individual ROI | AI, R, RT, AL, ML, CL, CM, RM, RTM, RTL, RTp, Tpt, CPB, RPB, STGr, TPO, TAa, area 45a, area 46d | Left, right and bilateral |
| Core family | AI, R, RT | Left, right and bilateral |
| Belt family | AL, ML, CL, CM, RM, RTM, RTL, RTp | Left, right and bilateral |
| Temporal/multisensory family | Tpt, CPB, RPB, STGr, TPO, TAa | Left, right and bilateral |
| Core-belt network | All Core and Belt ROIs | Bilateral |

**Supplementary Table 5. Decoding performance during the audiovisual task.** A) Three-condition balanced accuracy for auditory-only (AC), audiovisual-congruent (VAC) and audiovisual-incongruent (VAI) conditions in four a priori bilateral ROI families. CTR and OT estimates are equal-weight means across M50 and M58 and are reported with 95% run-level bootstrap confidence intervals. Permutation P and q values test performance against the 33.3% chance level and are reported to four decimal places. B) AC recall from the same three-condition classifier. Direct Injection effects were expressed as OT–CTR differences and assessed by exact reassignment of complete pharmacological sessions within each monkey, yielding 36 possible combined assignments across the two monkeys. Exact P values and FDR-adjusted q values are reported to four decimal places.

##### A. Three-condition balanced accuracy

| ROI family | CTR BA [95% CI] | CTR P; q | OT BA [95% CI] | OT P; q | OT–CTR ΔBA | Exact P; q |
|---|---|---|---|---|---|---|
| Auditory Core | 0.465 [0.419, 0.510] | P=.001; q=.001 | 0.499 [0.454, 0.538] | P=.001; q=.001 | 0.033 | P=.6111; q=.9028 |
| Auditory Belt | 0.489 [0.441, 0.536] | P=.001; q=.001 | 0.523 [0.478, 0.572] | P=.001; q=.001 | 0.034 | P=.7222; q=.9028 |

| | | | | | | |
|---|---|---|---|---|---|---|
| Temporal multisensory | 0.639 [0.587, 0.691] | P=.001; q=.001 | 0.641 [0.597, 0.685] | P=.001; q=.001 | 0.002 | P=1.0000; q=1.0000 |
| Frontal | 0.505 [0.460, 0.556] | P=.001; q=.001 | 0.482 [0.439, 0.523] | P=.001; q=.001 | -0.023 | P=.6667; q=.9028 |

**B. Auditory-only recall**

| ROI family | CTR AC recall | CTR P; q | OT AC recall | OT P; q | OT–CTR Δ recall | Exact P; q |
|---|---|---|---|---|---|---|
| Auditory Core | 0.579 | P=.001; q=.001 | 0.539 | P=.001; q=.001 | -0.039 | P=.8333; q=.8333 |
| Auditory Belt | 0.610 | P=.001; q=.001 | 0.690 | P=.001; q=.001 | 0.079 | P=.5000; q=.6667 |
| Temporal multisensory | 0.854 | P=.001; q=.001 | 0.872 | P=.001; q=.001 | 0.018 | P=.5000; q=.6667 |
| Frontal | 0.557 | P=.001; q=.001 | 0.627 | P=.001; q=.001 | 0.070 | P=.4444; q=.6667 |

**Supplementary Table 6. Condition-specific resting-state functional connectivity and OT-CTR differences across selected predefined ROI-group pairs.**

ROI groups were specified a priori from RetCHARM and SARM atlas regions according to anatomical and functional subdivisions and were not derived by clustering the resting-state connectivity data. For each ROI-group pair, Fisher-z-transformed connectivity was averaged across all constituent ROI-to-ROI edges within each run. Pooled estimates are equal-weight means across M50 left, M50 right, M58 left and M58 right and are reported with conditional 95% confidence intervals. CTR and OT connectivity were tested against zero using two-sided stratified sign-flip permutation tests. OT-CTR differences were tested using stratified run-label permutations, with Benjamini-Hochberg FDR correction across the predefined ROI-group comparisons. Same-direction agreement indicates the number of the four monkey-by-hemisphere combinations showing the same OT-CTR direction as the pooled effect; 4/4 therefore indicates concordant effects in both hemispheres of both monkeys. The M50 and M58 columns indicate whether the corresponding monkey-specific OT-CTR permutation test survived BH-FDR correction across the same predefined comparisons, with the two hemispheres given equal weight. Because each injection condition was represented by one resting-state session per monkey, monkey-specific results provide supporting within-dataset evidence rather than independent session-level replication.

| ROI-group pair | CTR Fisher z [95% CI]; q | OT Fisher z [95% CI]; q | OT-CTR Δz [95% CI]; Pperm; q | Same direction | M50 FDR q<.05 | M58 FDR q<.05 | Interpretation |
|---|---|---|---|---|---|---|---|
| Auditory Core-Belt | 0.220 [0.207, 0.234]; q=2.00×10⁻⁴ | 0.255 [0.244, 0.268]; q=3.14×10⁻⁴ | 0.036 [0.017, 0.054] **Pperm=.001; q=.006** | 4/4 | **Yes** | No | Stronger pre-existing positive connectivity |
| Auditory Core/Belt-early visual | -0.060 [-0.072, -0.048]; q=2.00×10⁻⁴ | -0.016 [-0.033, 0.000]; q=.126 | 0.043 [0.023, 0.065] **Pperm=2.00×10⁻⁴; q=.002** | 4/4 | **Yes** | No | Negative connectivity attenuated toward zero |
| Auditory Core/Belt-visual motion | 0.014 [0.004, 0.026]; q=.045 | 0.037 [0.025, 0.050]; q=3.14×10⁻⁴ | 0.023 [0.006, 0.040] **Pperm=.014; q=.029** | 3/4 | **Yes** | No | Stronger pre-existing positive connectivity |
| Auditory Core/Belt-extended visual | -0.013 [-0.021, -0.005]; q=.009 | 0.012 [-0.000, 0.024]; q=.130 | 0.025 [0.010, 0.039] **Pperm=.002; q=.008** | 4/4 | **Yes** | No | Negative connectivity attenuated toward zero; OT was not significantly positive |

| | | | | | | | |
|---|---|---|---|---|---|---|---|
| Auditory Core/Belt-VLPFC | -0.003 [-0.017, 0.012]; q=.780 | 0.032 [0.013, 0.050]; q=.004 | 0.034 [0.011, 0.057] **Pperm=.007; q=.020** | 4/4 | **Yes** | No | Significant positive connectivity emerged under OT |
| Auditory Core/Belt-dorsal/oculomotor PFC | -0.028 [-0.043, -0.014]; q=.001 | 0.003 [-0.011, 0.018]; q=.721 | 0.032 [0.011, 0.052] **Pperm=.006; q=.017** | 3/4 | **Yes** | No | Negative connectivity attenuated toward zero |
| Visual-mPFC | -0.119 [-0.134, -0.103]; q=2.00×10⁻⁴ | -0.081 [-0.103, -0.058]; q=3.14×10⁻⁴ | 0.038 [0.010, 0.065] **Pperm=.013; q=.029** | 3/4 | **Yes** | No | Negative under both treatments, but less negative under OT |
| Visual-amygdala | -0.037 [-0.048, -0.026]; q=2.00×10⁻⁴ | -0.012 [-0.022, -0.002]; q=.048 | 0.025 [0.011, 0.039] **Pperm=.003; q=.009** | 3/4 | No | **Yes** | Negative under both treatments, but less negative under OT |
| Auditory Core/Belt-claustrum/PAG | 0.040 [0.029, 0.051]; q=2.00×10⁻⁴ | 0.009 [-0.002, 0.020]; q=.244 | -0.031 [-0.047, -0.015] **Pperm=4.00×10⁻⁴; q=.003** | 4/4 | **Yes** | No | Significant positive connectivity was reduced toward zero |
| Auditory Core/Belt-dorsal premotor | 0.050 [0.032, 0.067]; q=2.00×10⁻⁴ | 0.006 [-0.012, 0.025]; q=.671 | -0.043 [-0.068, -0.019] **Pperm=.004; q=.014** | 3/4 | No | **Yes** | Significant positive connectivity was reduced toward zero |
| Insula-amygdala | 0.018 [0.005, 0.030]; q=.029 | -0.015 [-0.032, -0.000]; q=.130 | -0.032 [-0.054, -0.013] **Pperm=.004; q=.013** | 3/4 | **Yes** | No | Positive connectivity was lost; OT estimate was negative but nonsignificant |

Note. Rows are restricted to pooled OT-CTR effects reported in the main resting-state results and surviving pooled FDR correction. 'Yes' in the M50 or M58 column denotes monkey-specific BH-FDR significance; exact monkey-specific q values are not reproduced here.

**Supplementary Table 7. FDR-significant ROI-pair differences in across-run covariation of task-evoked PSC.**
Spearman correlations are shown descriptively under control (CTR) and oxytocin (OT). Δ Fisher z = z(OT) − z(CTR); positive values indicate greater/more positive covariation under OT. Only ROI-pair effects surviving FDR correction are shown.

**A. Unimodal auditory task**

| Connection family | ROI pair | ρ CTR | ρ OT | Δ Fisher z (OT−CTR) | p interaction | q FDR | Directional consistency |
|---|---|---|---|---|---|---|---|
| Auditory Core–VLPFC | RT–area 45a | 0.383 | -0.279 | -0.691 | $1.13 \times 10^{-4}$ | .0014 | 4/4 |
| Auditory Core–VLPFC | RT–area 46v | 0.353 | -0.171 | -0.542 | .0015 | .0089 | 4/4 |
| Auditory Core–mPFC | AI–area 9m | 0.300 | -0.292 | -0.611 | .0018 | .0107 | 3/4 |
| Auditory Core–mPFC | R–area 9m | 0.114 | -0.321 | -0.448 | .0039 | .0118 | 4/4 |
| Auditory Belt–mPFC | CL–area 10mr | 0.255 | -0.148 | -0.409 | .0046 | .0245 | 3/4 |
| Auditory Belt–mPFC | CM–area 9m | 0.294 | -0.201 | -0.507 | .0025 | .0199 | 4/4 |

| | | | | | | | |
|---|---|---|---|---|---|---|---|
| Auditory Belt–mPFC | RM–area 9m | 0.186 | -0.186 | -0.376 | .0011 | .0181 | 4/4 |
| Early visual–Dorsal/oculomotor PFC | V3d–area 8Av | 0.380 | -0.171 | -0.572 | $4.94 \times 10^{-4}$ | .0104 | 4/4 |
| Insula–Dorsal/oculomotor PFC | Ia/Id–area 8Bs | 0.224 | -0.108 | -0.336 | .0038 | .0344 | 4/4 |
| Dorsal/oculomotor PFC–Claustrum | area 8Bs–Cl | 0.182 | -0.173 | -0.359 | .0079 | .0236 | 4/4 |

**B. Auditory-only condition of the audiovisual task**

| Connection family | ROI pair | ρ CTR | ρ OT | Δ Fisher z (OT−CTR) | p interaction | q FDR | Directional consistency |
|---|---|---|---|---|---|---|---|
| Auditory Core–Early visual | R–V3d | -0.032 | 0.182 | 0.217 | $8.50 \times 10^{-4}$ | .0179 | 3/4 |
| Auditory Core–Insula | AI–Ri | 0.089 | 0.423 | 0.362 | .003 | .027 | 4/4 |
| Auditory Core–PAG | AI–PAG | 0.213 | -0.067 | -0.283 | .0116 | .0349 | 4/4 |
| Early visual–VLPFC | V3d–area 45a | 0.046 | 0.358 | 0.328 | .003 | .042 | 3/4 |
| Early visual–VLPFC | V3d–area 45b | 0.050 | 0.408 | 0.383 | .0013 | .0366 | 4/4 |
| Early visual–Claustrum | V1–Cl | -0.074 | 0.266 | 0.347 | .0129 | .0302 | 4/4 |
| Early visual–Claustrum | V2v–Cl | -0.149 | 0.228 | 0.383 | .002 | .0141 | 4/4 |
| Early visual–Claustrum | V3v–Cl | 0.007 | 0.304 | 0.308 | .0107 | .0302 | 4/4 |
| Insula–VLPFC | Ia/Id–area 45b | 0.043 | 0.290 | 0.256 | .0055 | .0329 | 4/4 |
| Insula–VLPFC | Ri–area 45b | -0.049 | 0.187 | 0.238 | .0048 | .0329 | 4/4 |
| VLPFC–Dorsal premotor | area_44–PMdr | 0.007 | 0.303 | 0.306 | .0048 | .0386 | 4/4 |
| VLPFC–PAG | area_44–PAG | -0.237 | 0.123 | 0.365 | .0131 | .0298 | 4/4 |
| VLPFC–PAG | area 45b–PAG | -0.106 | 0.234 | 0.344 | .0149 | .0298 | 4/4 |
| Dorsal/oculomotor PFC–Dorsal/oculomotor PFC | area 8Ad–area 8Bs | 0.410 | 0.610 | 0.273 | .0037 | .011 | 4/4 |
| Dorsal/oculomotor PFC–Dorsal premotor | area 8Ad–PMdr | 0.009 | 0.370 | 0.379 | $3.37 \times 10^{-4}$ | .002 | 4/4 |
| mPFC–PAG | area 9m–PAG | 0.217 | 0.003 | -0.218 | .0216 | .0433 | 4/4 |
| Claustrum–Amygdala | Cl–PaL | -0.145 | 0.230 | 0.381 | $9.71 \times 10^{-4}$ | .0058 | 4/4 |
| PAG–Amygdala | PAG–PaL | -0.100 | 0.166 | 0.268 | .0043 | .0257 | 4/4 |

Note. p values are from the Xz × Treatment interaction in rank-standardized run-level mixed-effects models. q values are Benjamini–Hochberg FDR-adjusted within the corresponding predefined connection-family block. Directional consistency is the number of monkey-by-hemisphere strata (M50-left, M50-right, M58-left, M58-right) showing the same OT−CTR direction as the pooled effect. This analysis quantifies across-run covariation of task-evoked response amplitudes and is not a measure of temporal functional connectivity.
